# A 180 My-old female-specific genome region in sturgeon reveals the oldest known vertebrate sex determining system with undifferentiated sex chromosomes

**DOI:** 10.1101/2020.10.10.334367

**Authors:** Heiner Kuhl, Yann Guiguen, Christin Höhne, Eva Kreuz, Kang Du, Christophe Klopp, Céline Lopez-Roques, Elena Santidrian Yebra-Pimentel, Mitica Ciorpac, Jörn Gessner, Daniela Holostenco, Wibke Kleiner, Klaus Kohlmann, Dunja K. Lamatsch, Dmitry Prokopov, Anastasia Bestin, Emmanuel Bonpunt, Bastien Debeuf, Pierrick Haffray, Romain Morvezen, Pierre Patrice, Radu Suciu, Ron Dirks, Sven Wuertz, Werner Kloas, Manfred Schartl, Matthias Stöck

## Abstract

Several hypotheses explain the prevalence of undifferentiated sex chromosomes in poikilothermic vertebrates. Turnovers change the master sex determination gene, the sex chromosome or the sex determination system (e.g. XY to WZ). Jumping master genes stay main triggers but translocate to other chromosomes. Occasional recombination (e.g. in sex-reversed females) prevents sex chromosome degeneration. Recent research has uncovered conserved heteromorphic or even homomorphic sex chromosomes in several clades of non-avian and non-mammalian vertebrates. Sex determination in sturgeons (Acipenseridae) has been a long-standing basic biological question, linked to economical demands by the caviar-producing aquaculture. Here, we report the discovery of a sex-specific sequence from sterlet (*Acipenser ruthenus*). Using chromosome-scale assemblies and pool-sequencing, we first identified a ~16 kb female-specific region. We developed a PCR-genotyping test, yielding female-specific products in six species, spanning the entire phylogeny with the most divergent extant lineages (*A. sturio, A. oxyrinchus* vs. *A. ruthenus, Huso huso*), stemming from an ancient tetraploidization. Similar results were obtained in two octoploid species (*A. gueldenstaedtii, A. baerii*). Conservation of a female-specific sequence for a long period, representing 180 My of sturgeon evolution, and across at least one polyploidization event, raises many interesting biological questions. We discuss a conserved undifferentiated sex chromosome system with a ZZ/ZW-mode of sex determination and potential alternatives.

## Introduction

Sexual reproduction is an old feature of life. In vertebrates, sexual development is determined by environmental triggers (Environmental Sex determination, ESD) or genotypic sex determination (GSD) (e.g.[1]) or a combination thereof. In an existing GSD-system, the paradigmatic evolutionary model of sex chromosome evolution suggests the rise of a new sex chromosome from an autosome by mutation or translocation of a gene, which becomes a trigger of the sex-determining regulatory networks. Heterozygotes encode one sex and homozygotes the other [2–4]. According to this classical view, mutations and/or linkage to genes with sexually antagonistic effects may be favoured near this gene, i.e. mainly in unpaired parts of Y and W chromosomes, profiting from linkage disequilibrium. To preserve epistasis, recombination between such mutations and the sex-determining locus would then be suppressed in the heterogametic sex [5–6]. Such nonrecombining sex chromosomes are expected to degenerate through the progressive accumulation of deleterious mutations [7–8], except for small pseudo-autosomal regions. Recently, as an alternative or additional cause of degeneration, instability and divergence of *cis*-regulatory sequences in non-recombining genome regions, which become selectively haploidised to mask deleterious mutations on coding sequences, have been suggested [9].

In contrast to the conservation of differentiated sex chromosomes in most mammals and birds, in the majority of poikilothermic vertebrates differentiated sex chromosomes are rarely observed [10–12] (but see [13]). Several hypotheses have been offered to explain the prevalence of undifferentiated sex chromosomes. The high-turnover model proposes that the emergence of new master sex-determining genes on new chromosomes would prevent sex chromosome degeneration [14–17]. The “jumping sex locus”-hypothesis [18] implies that the sex determination gene is maintained but changes its position to different chromosomes, preventing evolutionary decay of the original sex chromosome [19]. The ‘fountain of youth’ hypothesis [20] suggests that sex chromosomes may escape from degeneration by occasional recombination Because sex-specific recombination depends on phenotypic rather than genotypic sex, homomorphic X and Y chromosomes might recombine in sex-reversed females. These rare events should generate bursts of new Y haplotypes, which will be quickly sorted out by natural or sexual selection [20]. Seemingly young sex chromosomes may thus carry old-established sex-determining genes, challenging the opinion that sex chromosomes unavoidably decay [21–22].

Beyond mammals and birds, conserved sex chromosomes have recently been discovered in several amniote (specifically reptile) clades [23–26], all of which feature differentiated sex chromosomes. Evolutionarily very old, conserved and homomorphic ZZ/ZW sex chromosomes are known in some ratite birds (Ratidae), dating back >130 My [27–28]. Similarly, skinks (Scincidae) share homologous, mostly poorly differentiated XX/XY sex chromosomes across a wide phylogenetic spectrum for at least 85 million years [29]. Recent findings in the teleost family Esocidae report undifferentiated sex chromosomes of similar evolutionary age (65-90 My) in teleosts [30]. The Salmonidae family, which experienced a whole genome duplication ca. 90 Mya [31], presumably harbours a conserved sex determination gene, perhaps as old as 50 My, on different chromosomes in different species [18–19], which encodes a homomorphic XX/XY sex chromosome system. Here we report the discovery of a sex-specific region in sturgeon (Acipenseridae), which is preserved in many extant species with a common ancestor about 180 Mya.

Sturgeons and paddlefish comprise 27 living species [32], which branched off 330 Mya [33], from the root of the more than 31,000 living teleosts. The evolutionary history of vertebrates is characterized by several rounds of whole genome duplications (WGD). The 1R and 2R WGD happened early on, before the jawed vertebrates appeared, but the sturgeon branch diverged before the teleost 3R WGD [34–35]. Later on, sturgeon experienced several clade-specific polyploidizations, resulting in chromosome numbers from ~120 up to ~380 [36]. Based on a chromosome-scale genome assembly of *Acipenser ruthenus*, Du *et al.* [33] have recently shown a WGS, early in the sturgeon radiation. Although subsequent rediploidization of the genome caused the loss of entire chromosomes or large fragments (i.e. segmental deduplication), structural and functional tetraploidy was maintained over 180 million years, and a decrease of redundancy in the tetraploid genome follows mostly random processes. Sturgeons possess undifferentiated microscopically indistinguishable sex chromosomes [33,37]. Although the sex ratio of offspring from experimental gynogenesis led to the assumption that all sturgeons possess female heterogametic (ZZ/ZW) sex chromosome systems [38], a sex-specific marker, confirming female heterogamety and allowing genetic studies, has not been found [33].

## Materials and Methods

### Animal sampling and establishment of genetic families

#### Acipenser ruthenus

For pool-sequencing (pool-seq) 61 (31 female, 30 male) adult sterlets from the broodstock at the Leibniz-IGB (Berlin, Germany), representing the Danube River population [33], were sexed by gonadal biopsies, evidence from reproduction and a few autopsies. In addition, 52 sterlets (table 1) were sampled from three aquaculture populations. These came from two other *A. ruthenus* brood stocks, originating from the Danube (Wöllershof, Bavaria, Germany, 20-25 cm, ca. 1 year old; and Beucha, Saxony, Germany, 10-12 cm, 6 months old), as well as *A. ruthenus* interpopulation hybrids from the Volga and Danube Rivers (12-20 cm, 8-10 months). They were euthanized and gonadal sex determined histologically, and finclips used for genotyping. In spring 2020, a genetic family was derived from the IGB-broodstock; at the age of 4 months, histological sections of gonads from 14 offspring were prepared, and finclips from them and their parents also used for genetics.

**Table 1.** Numbers of individuals tested with the sex-linked PCR marker *AllWSex2* (p-values for association with sex were calculated using a Pearson’s Chi-squared test with Yates’ continuity correction; gel pictures in electronic supplementary material, figure S3–S10). To avoid circular evidence, neither the 61 (30 females, 31 males) *A. ruthenus* used for pool-sequencing, nor the offspring of the three genetic families of *A. ruthenus, A. oxyrhinchus* and *H. huso* (supplementary materials, figures S8–S10) but only their parents are counted here.

| Species | Ploidy | Positive Females | Positive Males | *p*-value of association with sex | Type of phenotypic sexing |
| --- | --- | --- | --- | --- | --- |
| *A. sturio* | 4n | 18/18 | 0/18 | 1.456e-08 | ultrasound, biopsy if contradiction to genotype, few autopsies |
| *A. oxyrinchus* | 4n | 12/12 | 0/15 | 1.536e-06 | adult breeders with known reproduction, histology in family offspring |
| *H. huso* | 4n | 12/12 | 0/16 | 9.311e-07 | adult breeders with known reproduction, biopsy, histology in family offspring |
| *A. ruthenus* | 4n | 25/25 | 0/27 | 4.11e-12 | adult breeders with known reproduction, biopsy, histology in family offspring |
| *A. baerii* | 8n | 28/28 | 0/28 | 5.352e-13 | adult breeders with known production of gametes |
| *A. gueldenstaedtii* | 8n | 15/15 | 0/12 | 1.536e-06 | adult breeders with known reproduction, ultrasound, few autopsies |

#### Huso huso

Using minimal invasive biopsy samples from the gonads of 28 subadult and adult beluga sturgeon from two fish farms (Horia and Peceneaga, Romania, fish between 10 and 13 years old) were inspected by the unaided eye and/or microscopically after standard hematoxylin/eosin staining (described below). Finclips were sampled for genetic analyses (table 1). To generate genetic families, on April 6, 2018, a beluga sturgeon male (PIT-tag 000004691858; 200 cm, ~100 kg) was captured in the Danube delta region at river-Km 126 (distance from the Black Sea) in Romania. On April 13, 2018, a mature *H. huso* female (PIT 642099000568452; TL 260 cm, ~150 kg, near Km 100), and another male on April, 14, 2018 (PIT 968000004692054; TL 143 cm, ~22 kg, Km 126) were caught. All three were transferred to an aquaculture company (Horia, Romania). After stimulation of all the three wild-caught animals and one additional captive male with gonadotropin-releasing hormone (GnRH), ovulation started on April 16. Eggs were fertilized with sperm of each male separately. Fin clips of the four adult fish were taken, and the offspring of the individual matings were reared separately until the age of one year. All offspring fish were individually tagged with PIT-tags and then reared in common tanks. For this study, fish of one genetic family were sampled at the age of 15-24 months for gonadal histology and finclips. Wild-caught adults and, after two years, hundreds of their tagged offspring were released into the Danube to minimize the impact on this highly threatened species.

#### A. oxyrinchus

Tissue samples of 27 adult Baltic sturgeon (table 1) from the broodstock in Mecklenburg-Pomerania (Born, Germany), kept for re-introduction into Oder River and Baltic Sea, were taken from tagged males and females, sexed using gonadal biopsies and evidence from reproduction. In 2017, genetic families were derived from breeding pairs during the yearly reproduction. Finclips from the parental animals were kept in ethanol. A subset of juveniles was raised in separate tanks at the Leibniz-IGB (Berlin, Germany) for three years.

#### A. sturio

Tissue samples of 36 adult European sturgeon (table 1) from a broodstock of immature fish at the IGB, raised for conservation aquaculture to re-introduce the species into the Elbe River were initially sexed using ultrasonic evidence. Gonadal biopsies were taken if genotyping did not match the ultrasonic evidence. Few specimens came from autopsy of deceased fish.

#### A. baerii

Finclips of 56 adult Siberian sturgeon (table 1) used as breeders with known phenotypic sex were sampled in an aquaculture breeding company (l’Esturgeonnière, France) and kept in 100% ethanol until DNA extraction.

#### A. gueldenstaedtii

Finclips of 27 adult Russian sturgeons (table 1) were sampled in an aquaculture breeding company (Ecloserie de Guyenne, SCEA Sturgeon, France) and kept in 100% ethanol until DNA extraction. All fish were sexed based on evidence from ultrasound, reproduction and autopsy.

### DNA processing for genotyping

DNA from finclips or other tissue samples was extracted using the DNeasy Tissue Kit (Qiagen) or the BioSprint robotic workstation with the 96 DNA Plant Kit (Qiagen, Germany), according to the manufacturer’s protocols.

### Histological sexing of juvenile and subadult fish as well as offspring of genetic families

Offspring from genetic families of *H. huso, A. ruthenus*, and *A. oxyrinchus* was sacrificed by an overdose of buffered tricaine methanesulfonate (MS222; 0.3 g/L, PHARMAQ), individual length and wet weight were recorded, finclips (stored in 100% ethanol) and gonadal samples were taken immediately. The right gonad strand was fixed in phosphate-buffered formaldehyde solution (ROTI®Histofix, 4%, Carl Roth) for at least 24 h at room temperature, then rinsed three times for 24 h with 70% ethanol, and stored at room temperature. Gonad strands were dehydrated in an increasing series of ethanol (75%, 90%, 95%, 100%), rinsed in xylol (Carl Roth), and transferred into Paraplast® (Leica), using the Shandon Excelsior ES Tissue Processor (Thermo Fisher Scientific). Gonads were embedded in a spiral-like arrangement, to ideally cut the whole organ into 5 µm sections across its entire length (microtomes: Leica 2065 or Microm HM 340E, Thermo Fisher Scientific). Sections were mounted on glass slides, and stained using standard hematoxylin/eosin (0.1%, Carl Roth). Histological evaluation was made at various magnifications using light microscope Nikon Eclipse Ni-U with Nikon DS-Fi3 camera, and the corresponding software Nikon DS-L4 to archive images.

### Pool-sequencing

The genomic DNA of 31 female and 30 male *A. ruthenus* specimens was extracted from 90% ethanol-preserved fin clips using a classical phenol/chloroform protocol, quantified using Qubit fluorimetry and analysed using the Fragment Analyzer (Advanced Analytical Technologies, Inc., Iowa, USA). DNA shorts read sequencing was performed at the GeT-PlaGe core facility, INRAe Toulouse (http://www.get.genotoul.fr). Two DNA pool-seq libraries were prepared according to manufacturer’s protocols using the Illumina TruSeq Nano DNA HT Library Prep Kit (Illumina, California, USA). Briefly, from 200 ng of each sample, DNA was fragmented (550 bp) by sonication on a M220 Focused-ultrasonicator (COVARIS). Size selection was performed using SPB beads (kit beads) and 3’-ends of the blunt fragments were mono-adenylated. Then, adaptors and indexes were ligated and the construction amplified with Illumina-specific primers (8 cycles). Library-quality was assessed using Fragment Analyzer and libraries were quantified by qPCR using the Kapa Library Quantification Kit (Roche). Sequencing was performed on a NovaSeq S4 lane (Illumina, California, USA), using a paired-end read length of 2×150 bp with the Illumina NovaSeq Reagent Kits.

### Analysis of sex specific variants based on a male reference genome

Pool-seq data were mapped to the male *A. ruthenus* reference genome (*Acipenser_ruthenus*.ARUT1.2.dna.toplevel.fa, which is nearly identical to the current reference genome at NCBI GCF_010645085.1; electronic supplementary material, table S1) by Minimap2 (with parameters –t 12 –x sr –a; [39]) and converted to sorted bam-files by samtools [40]. Variants (SNPs, MNPs, INDELs) were called from both bam files using the Platypus variant caller [41] with integrated duplicate fragment reads removal (--filterDuplicates=1) and reassembly of reads (--assembleAll=1). The resulting vcf-file was screened for sex-specific variants using awk-scripting and different variant read coverage cutoffs (i.e. maximum reads in males (0, 1 …), minimum reads in females (10, 13 …) or *vice versa*). Variants were clustered according to their distance in the genome (maximal distance of variants in a cluster: 4000 bp), counted using bedtools (merge, annotate; [42]) and sorted by variant count. Regions with highest variant densities and neighbouring regions were inspected for genes related to sex determination.

### Reconstruction of male and female specific genomic regions from unmapped pool-seq data

Unmapped reads potentially representing regions not present in the reference genome were extracted from bam-files and assembled using the IDBA assembler [43]. The resulting male/female specific contigs were added to the reference genome and pool-seq data was mapped again (as described above) to this extended reference genome. Finally, sequencing coverage by female and male pools was compared in bedgraph files derived from the bam-files (bedtools genomecov –bga –split – ibam …; bedtools unionbedgraph). These were screened for putative haploid coverage in one sex and no or low, spurious coverage in the other sex to identify female/male specific regions.

Another high-quality assembly of a female *A. ruthenus* has recently been published at NCBI by the Vertebrate Genomes Project (GCA_902713425.1). We repeated our analyses from above using this reference with an emphasis on detection of coverage differences. Using bedgraph-files we compared female and male pool-sequencing coverage at the bp-level along the genome. We used different filtering approaches to call regions which had no or low coverage in one sex and approximately haploid coverage in the other sex. The most successful filtering applied awk-scripting to filter regions in male and female genomes where 0 coverage in male pool and a coverage of larger than 0.3 in the female pool was observed (here 1.0 was the normalized diploid coverage). These regions were combined in larger blocks, if they were as close as 30 bp to each other (bedtools merge). Finally, we filtered for blocks that were larger than 160 bp and derived at a clear signal present only in female data and female genome.

### Primer design and PCR conditions

Using the male *vs*. female *A. ruthenus* genome sequence as well as preliminary Illumina-data of a female *A. oxyrinchus* genome, we designed a number of primers, which were primarily tested in gradient-PCRs on phenotypically sexed *A. ruthenus*. While several primers showed sex-specific amplification in this species, one (*AllWSex2*) could be optimized to work across multiple species (primer sequences and PCR-conditions: electronic supplementary material, texts S1-S3).

### Phylogenomic reconstruction

To build a phylogenetic tree for sturgeons, we download transcriptome assemblies of *Acipenser baerii*, *A. oxyrinchus*, *A. schrencki*, *A. sinensis* and *A. transmontanus* from the Public Sturgeon Transcripts Database (http://publicsturgeon.sigenae.org/home.html) and of *Acipenser gueldenstaedtii* from NCBI (https://www.ncbi.nlm.nih.gov/Traces/wgs/?page=1&view=tsa&search=acipenser from NCBI (https://www.ncbi.nlm.nih.gov/Traces/wgs/?page=1&view=tsa&search=acipenser). The transcriptome of *Huso huso* was *de novo* assembled using Trinity [44] from mixed organs. Annotated genomes of *A. ruthenus* and *Latimeria chalumnae* were also downloaded from NCBI (electronic supplementary material, table S2). To identify orthologs between two species, we used the “Reciprocal best hit (RBH)” method, and *A. ruthenus* as the hub to retrieve orthologous genes across all species. Protein sequences of the transcriptome contigs were predicted using GeneWise [45] by aligning each contig to its orthologous protein in *A. ruthenus*. Contigs with less than 60% of the protein aligned were discarded. In total, we collected 1017 genes with orthology across all species. For each orthologous gene, the protein sequences were aligned across all species using MAFFT [46], and trimmed them, using trimAl [47]. All 1017 alignments were then combined in a concatenated alignment, consisting of 383506 aligned amino acid sites. Finally, this alignment was transferred to RAxML [48] to construct a Maximum Likelihood-phylogeny.

## Results

### Analyses of sterlet (A. ruthenus) female and male pool-sequencing

Pool-sequencing resulted in 553,117,774 reads (83.5 Gbp) and 486,762,084 reads (73.5 Gb) for the female and the male *A. ruthenus* pools, respectively. Pool‐sequencing Illumina reads are available in the Sequence Read Archive (SRA: SRX9341540; SRX9341541), under BioProject reference PRJNA670610. After mapping to the reference genomes, the peak of the unimodal coverage distribution was 42x for the female pool and 37x for the male pool. There was no significant difference in the percentage of mapped reads for both pools (96.9% reads mapped).

We initially used two bioinformatic strategies to uncover female-specific pool-seq data in *A. ruthenus*. We first analysed sex-specific variants based on a male reference genome. The number of female or male specific variants was low, but their ratio already hinted at a ZW sex determination system (details: electronic supplementary material, text S4). In the top candidate region, which harboured 27 female-specific variants at HiC_scaffold_4:45,149,338-45,165,057 (electronic supplementary material, figure S1, table S1, text S5), we found a female-specific 15 bp-indel, which we used to design PCR-primers. These amplified a product concordant with phenotypic sex of some *A. ruthenus*, but not all (data not shown). We also analysed male- or female-specific pool-seq reads, not matching the male reference genome, and independently assembled these unmapped reads from the male/female pools. This resulted in 1,067 additional female-specific contigs spanning 658,732 bp. However, BlastX-searches using these contigs did not detect any sequences related to known sex determination genes.

### Identification of a female-specific locus in A. ruthenus by coverage analysis using pool-seq data and a female reference genome

Due to a possible high repeat content of a W-locus, short-read assemblies might not be able to resolve it in large contigs but in rather fragmented small contigs, hindering further analysis. Recently, a long-read female haplotype reference of *A. ruthenus* has become available (GCA_902713425.1), which allowed remapping our pool-seq data and focusing on regions with different coverage between males and females. With stringent filtering parameters (no coverage in males; a minimum of 30% diploid coverage in females and minimum size of regions of 160 bp), we found 11 neighbouring matches (summing up to 2705 bp) in the female genome, while we did only find a single spurious match (185 bp) in the male genome. *Vice versa*, filtering for zero coverage in females and 30% of diploid coverage in males, also resulted in no hits (female genome) or two spurious hits (175 and 181 bp; male genome)

The filtered regions in the female genome fell into the region CACTIG010000179.1:61,229,779-61,285,849 (including 20 kb up- and down-stream). This region corresponds to the region HiC_scaffold_4:43,938,061-43,987,561 in the male assembly as shown by the alignments of the 20 Kbp up- and down-stream sequences. Figure 1 illustrates the female and the male pool-seq coverage of the identified locus. The female specific locus is located 1.162 Mb upstream of the identified cluster of 27 female-specific variants, mentioned above. Furthermore, a single contig (559 bp) of the 1,067 female-specific contigs assembled from the female pool-seq data, did match the identified locus. Thus, three independent analyses of the pool-seq data, specifically sequence variants, assembled female-specific reads and differential sequence coverage of the female genome assembly, support HiC_scaffold_4 as a sex chromosomal sequence, albeit with different specificity and resolution.

**Figure 1.**
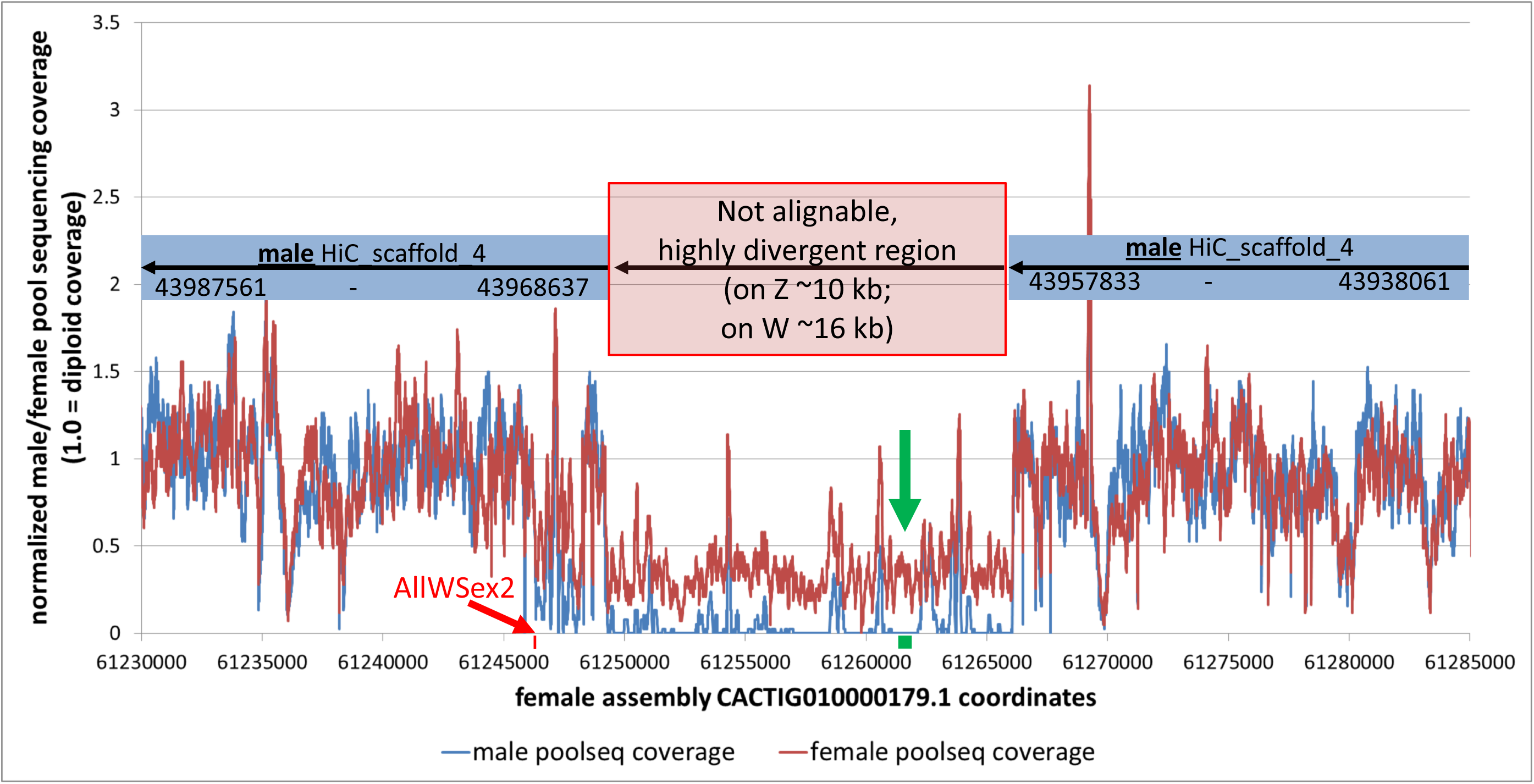
Detection of a female-specific locus in *Acipenser ruthenus*, when mapping female/male pool-sequencing reads to a female (ZW) reference genome assembly. Genome-wide signal detection was highly specific, when screening for larger regions (here >= 160 bp) with zero and non-zero coverage between sexes. The red block and arrow show the position of the *AllWSex2*-PCR-marker (CACTIG010000179.1: 61246236 - 61246344), which distinguishes sex in six sturgeon species. The green block and arrow mark the location of the only contig matching the region (CACTIG010000179.1: 61261166 - 61261703), when using the female-specific pool-seq read assembly approach. For final submissions, figures should be uploaded as separate, high resolution, figure files.

### Establishment of a W-linked PCR across the sturgeon species tree

Our PCR-marker *AllWSex2* revealed female-specific products for all unambiguously phenotypically sexed females under identical PCR-conditions in six sturgeon species (table 1), spanning the entire phylogenetic tree and the most divergent extant lineages (*A. sturio, A. oxyrinchus* vs. *A. ruthenus, H. huso*), all of which are tetraploid (~120 chromosomes). The marker showed also sex-linkage in two octoploid species (*A. gueldenstaedtii* and *A. baerii*; table 1, figure 2 and figure 3).

**Figure 2.**
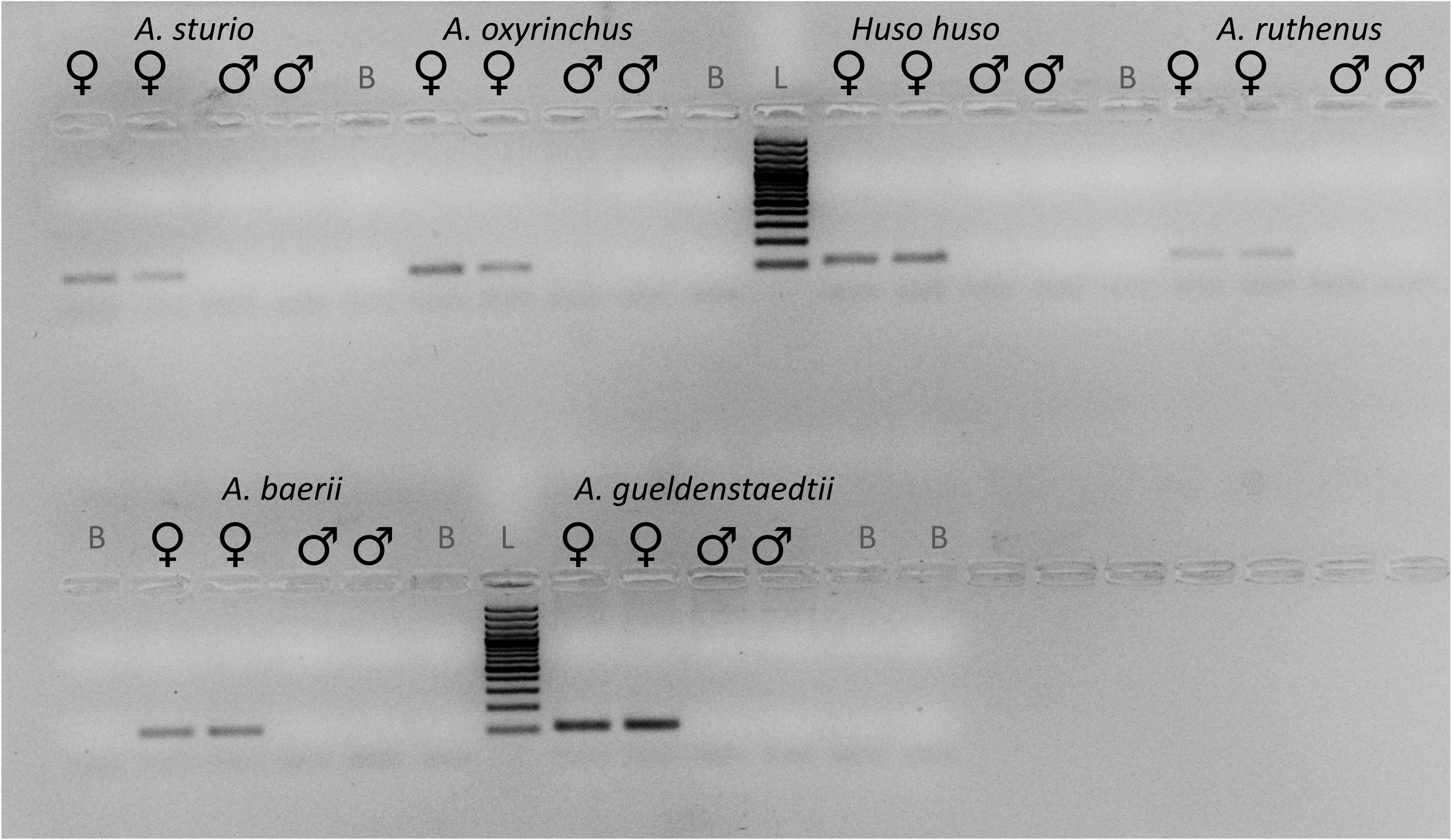
PCR products using the *AllWSex2* marker in six species across the sturgeon phylogeny under identical PCR-conditions (electronic supplementary material, texts S1 - S3) for two females and two males of each species. PCR products migrated ca. 45 min at 70 V on a 2% agarose gel in 1x TAE-buffer. Symbols and abbreviations: ♀: female; ♂: male; B: Blank, i.e. no DNA template in PCR; L: ladder, 100 bp-size-marker (Quantitas, Biozym).

**Figure 3.**
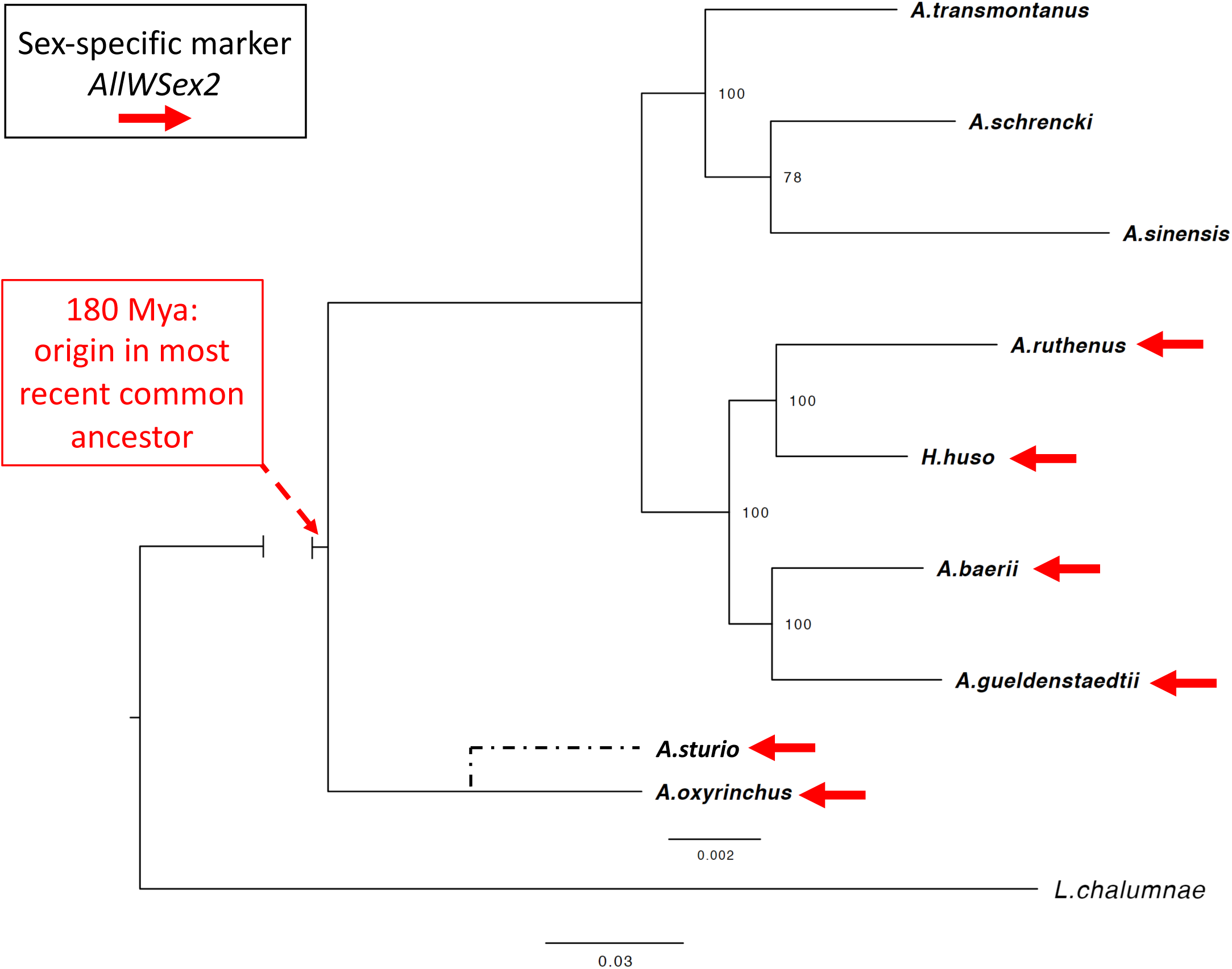
Phylogenomic tree of sturgeon species built by using RAxML with Maximum Likelihood (details: Materials and Methods). Numbers at nodes refer to bootstrap support values. The bar above indicates branch length of sturgeon lineages, while the bar below indicates branch length of the outgroup lineage coelacanth (*Latimeria chalumnae*). The approximate position of *A. sturio*, for which reliable transcriptomic data were unavailable, was added according to www.timetree.org. The root of Acipenseridae was dated at ~180 Mya, according to [33]. Red arrows mark those species, for we have obtained female-specific PCR-products, using primers *AllWSex2*_F/_R.

### Heredity of the sex-linked marker in three sturgeon species

In three genetic families, consisting of mother, father and their histologically sexed offspring, of *A. oxyrinchus*, *A. ruthenus* and *H. huso*, representing 180 My of sturgeon evolution, the marker *AllWSex2* was exclusively present in females, demonstrating mother to daughter transmission (gel pictures and histology in electronic supplementary material, figure S8 to figure S16).

## Discussion

We have discovered a female-specific (W) region, consistent with a potential ZZ/ZW system in sturgeon. This finding is in accordance with previous assumptions or indirect evidence that sturgeons exhibit a female-heterogametic sex determination system [37,49–50]. This genomic region is conserved in at least six sturgeon species, spanning the entire phylogenetic tree, represented by the most divergent species *A. sturio, A. oxyrinchus A. ruthenus*, and *H. huso*, all of which are formally ancient tetraploid (~120 chromosomes) [33]. In the *A. ruthenus* genome, the sequence is residing on chromosome 4 [33]. Of note, this genomic region is also sex-linked in two octoploid species (*A. gueldenstaedtii* with 258±4 chromosomes, and *A. baerii* with 249±5 chromosomes), demonstrating maintenance of this region and apparently a female heterogametic sex determination system through at least one round of polyploidization. In the future, conservation and evolution of the region in the remaining extant sturgeons should be examined to evaluate the degree to which this ancient female-specific genomic region is also conserved in other species. In particular, identification of the master sex determining gene(s) or other regulatory elements are required. At the current state of research, we also cannot rule out a situation like in Salmonidae, where a “jumping sex locus”, perhaps as old as 90 My [31], resides on different chromosomes in different species, all of which share a male heterogametic (XX/XY) system [18–19].

Our data reveal the oldest known vertebrate sex determining system with undifferentiated sex chromosomes. Conservation of a female-specific sequence for 180 My of sturgeon evolution, and across at least one additional polyploidization event, raises many interesting biological and evolutionary questions. We hypothesize that this region might be part of the sex locus. Given the slow protein evolution of the sturgeon genome, evolving distinctly slower than in teleosts, including basal species such as arowana and arapaima, but also slower than gar, coelacanth or elephant shark [33], we speculate that the relatively conserved karyotypes of the ancient tetraploid species examined here (*A. sturio, A. oxyrinchus*, *A. ruthenus, H. huso*) may have even maintained the same undifferentiated sex chromosome for 180 My. It will be fascinating to examine the mechanisms, which stabilized the sex determination system and whether or not this involves undifferentiated homologues sex chromosomes. Results from previous RADseq [33] and pool-sequencing of the same *A. ruthenus* (sampled from aquaculture populations, and with few if any family-effects expected), revealing 99.98 % of the chromosome as largely undifferentiated, let us assume that the sturgeon sex chromosome contains a large pseudo-autosomal region (PAR; [51–52]; electronic supplementary material, figure S1). This and the large chromosome number (~120 to ~260 for the 6 species) make it challenging to apply SNPs or microsatellites to examine whether *AllWSex2* resides on the same or different linkage groups and therefore is beyond the framework of this study, despite we have generated genetic families from three species. The disruption of sex determination in gonochoristic animals after genome duplication is considered the main cause for the relatively rare occurrence of polyploid animals compared to plants [53–54]. In vertebrates, in which autotetraploidy may be rarer than (hybrid-origin) allopolyploidy [55], polyploidy occurs especially frequently in amphibians [56], with few of which having their mostly undifferentiated sex chromosome characterized. In clawed frogs (*Xenopus*), alloploidy ranges from diploid to do-decaploid (12n) and subgenome evolution in allopolyploids has only recently been studied [57–58]. In *X. laevis*, the female-determining gene *DM-W* is a paralog of *DMRT1* [59–60], and arose after (and perhaps in response to) tetraploidization [61–63]. However, it is also found in some related *Xenopus* [61–62] but not in the entire radiation [64–65]. The clade-wide conservation of a male heterogametic sex determination gene and system (but not chromosome) in the 50 My-old, ancient tetraploid family Salmonidae (see above) curiously also evolved in a system with a common ancestral polyploidization event, as suggested here for Acipenseridae, with apparently conserved female heterogamety.

Our finding is too recent to allow comprehensive interpretation. Whole genome approaches are underway to address several of the most pressing questions, such as conservation of the region or the entire sex chromosomes in sturgeons. Conservation of the latter would even stronger challenge the classical paradigm of sex chromosome evolution [66–68] than the 180 My-long conservation of this small sex-linked sequence.

### Reasons for the challenges to identify a female-specific genomic region in sturgeon

The search for a sex chromosome or sex-linked markers in sturgeons has not been successful for a long time [37,49–50]. We discuss four major challenges, why this has been the case.

First, in *A. ruthenus*, the identified W-specific sequence is very short and resides on a large undifferentiated chromosome. It comprises only ~16 kb, corresponding to ~0.001% of the genome or ~0.02% of the corresponding chromosome (~106 Mb). It is impossible to detect such a small structural difference by karyotype imaging methods unless the W sequence is known and FISH probes can be designed.

Secondly, the high repeat content of the W-specific locus (~43%; www.repeatmasker.org) complicates its assembly by short-read data alone. Thus, when reconstructing female-specific genomic regions from unmapped pool-seq data, we filtered female-specific reads and independently assembled them. Unfortunately, the resulting small contigs were of limited utility for female-specific primer design. Looking for an explanation, we mapped these contigs back to the female genome assembly and found that only 0.09% of the sequence (a single contig) matched the W region, which we have identified by a sequencing coverage analysis.

Thirdly, short-read-based methods that call and analyse variants between female and male sequencing data suffered from the low number of sex-specific variants in sturgeons (few hundreds to thousands depending on filter parameters), especially when reduced representation methods like RADseq are used [33]. Even our whole genome pool-sequencing approach was not able to define the W-specific region by SNP markers. This approach only uncovered a nearby region (distance 1.16 Mb, ~1% of chromosome length); a result, which could point to reduced recombination around the W-specific sequence (W-linked SNPs). Finally, analysis of sequencing coverage enabled identification of the W-specific locus in *A. ruthenus*, but detection of haploid/diploid coverage differences were still complicated due to high variability (electronic supplementary material, figure S2), causing noisy signals for haploid genomic regions. Completely deleted regions in one sex compared to the other provided significant signals, but rely on the availability of a high-quality genome assembly of the heterogametic sex (figure 1).

### Potential application of the W-specific marker for conservation measures, aquaculture production and ethical issues

Beyond the basic biological interest in this ancient sex determination mechanism, molecular sexing of sturgeons is of great relevance in studies on wild sturgeon populations, their conservation and conservation aquaculture as well as caviar-producing aquaculture. Currently, ultrasonic diagnosis or biopsies are used for sexing. For ultrasonic sexing to become reliable, sturgeons have to reach progressed stages of maturation, six to ten years in some species, while biopsies are invasive and stress or even harm the fish [69]. Our molecular marker’s reliability outperforms methods like early biopsies or ultrasound-sexing. Future application of cotton skin swabs will make molecular sexing much less stressful for the fish. It will also aid conservation by reducing time and efforts to rear fish intended as future broodstock in *ex situ*-programs, reducing costs and improving the selection of candidates for living gene banks, not only based on rare alleles but also on sex. In this case, fish not selected for broodstock development can be released in the frame of recovery programs. In addition, in the caviar aquaculture, females, which are commercially more attractive, can be selected early on, and differentiated rearing for caviar (females) and meat (males) production becomes feasible. Releases of commercially reared fish are no option in conservation programs since they show restricted fitness due to intensive aquaculture [70]. In any case, we strongly emphasize that the early selection of females must not lead to littering male sturgeons, since their meat is a valuable source of protein [71].

## Permits and Licences

Permits to catch and propagate wild *H. huso* were provided by the Romanian Ministry of Agriculture and Rural Development (No. 7/19.03.2018). The license to keep and propagate sturgeon at the IGB is no. ZH 114 (LaGeSo, 18/02/2019).

## Acknowledgments

This work was funded through COFASP/ERANET (STURGEoNOMICS) by the German Federal Ministry of Food and Agriculture through the Federal Office for Agriculture and Food (grant nos. 2816ERA04G, 2816ERA05G) to M. Stöck, S.W., J.G. and M. Schartl; by the French National Research Agency (grant nos. ANR-16-COFA-0002) to YG, CK and CR, and by the Romanian Executive Agency for Higher Education, Research Development and Innovation Funding (UEFISCDI, contract 3/2017) to RS, MC and DH. This research was also in part supported by the Russian Science Foundation (RSF) to DP (grant number 18-44-04-007), by the Deutsche Forschungsgemeinschaft to MSch (DFG SCHA 408/14-1), and by a European Maritime and Fisheries Fund to YG (EMFF/FEAMP project SIBERSEX). Research on *A*. *gueldenstaedtii* was supported by a European Maritime and Fisheries Fund to YG (EMFF/FEAMP project S’STURGEON). We thank Marilena Maereanu, Katarina Tosic, and Marian Paraschiv for help with *H. huso* catching, propagation and raising of animals; Asja Vogt, Marcus Ebert, Jan Hallermann, Kevin Bäumler, Udo Wolf for help with *A. oxyrinchus* or *A. ruthenus* propagation and raising of animals; Elodie Dupin-de-Beyssat, Camille Eché, Antje Tillack, Sophia Lambert, Alina Heide, and Petra Kersten for help in the lab, Lukáš Kratochvíl for references, Jörg Plötner and Robert Schreiber for access to the Qiagen robotic workstation, and two anonymous reviewers for their constructive comments on a previous version of this paper..

## Electronic Supplementary material

Submitted in separate file: texts S1-S5, tables S1–S2, and figures S1–S16.

## Additional Information

### Ethics

The research using sturgeon in France, Romania and Germany were carried out in accordance with approved guidelines and obtained ethical approval by the institutional animal care and use committees; permits and licences as shown above.

### Data Accessibility

The pool‐sequencing Illumina reads are available in the Sequence Read Archive (SRA: SRX9341540 and SRX9341541), under BioProject reference PRJNA670610; all other datasets supporting this article have been uploaded as part of the Supplementary Material.

### Authors’ Contributions

Designed the study: MSchartl, MStöck, YG, SW, JG; wrote manuscript draft: HK, MS, MSch, YG, CH, JG, CK; analysed data and performed molecular analyses: HK, MStöck, CH, EK, CK, CLR, KD, ESYP, RD, JG, DP, WKleiner, KK, DKL; provided biological samples: RS, DH, MC, JG, HK, MStöck, WKloas, AB, EB, BD, PH, RM, PP. All authors read, contributed to and approved the manuscript.

## Electronic Supplementary Material

(Texts S1-S5, tables S1–S2, and figures S1–S16: pp. 1-10)

### Supplementary Texts

**Text S1.** *Primer sequences for marker* AllWSex2

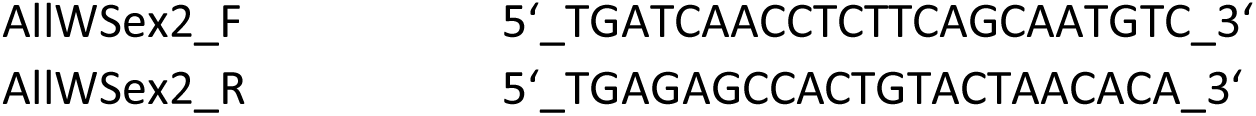

**Text S2.** *Pipetting scheme for AllWSex2-PCR*

|  | Final concentration | Stock | Volume [μl] |
| --- | --- | --- | --- |
| ddH<sub>2</sub>O |  |  | 17.3 |
| 10x TopTaqbuffer (Qiagen) | 1 | 10x | 2.5 |
| dNTP | 0.2 | [10mM] | 0.5 |
| AllWSex2_F | 0.4 | [10 pmol/μl] | 1.25 |
| AllWSex2_R | 0.4 | [10 pmol/μl] | 1.25 |
| TopTaq Polymerase (Qiagen) | 2 U/rxn | [120 rxn/24μl] | 0.2 |
| DNA | 20 ng/μl |  | 2.0 |

**Text S3.** *Temperature scheme for AllWSex2-PCR*

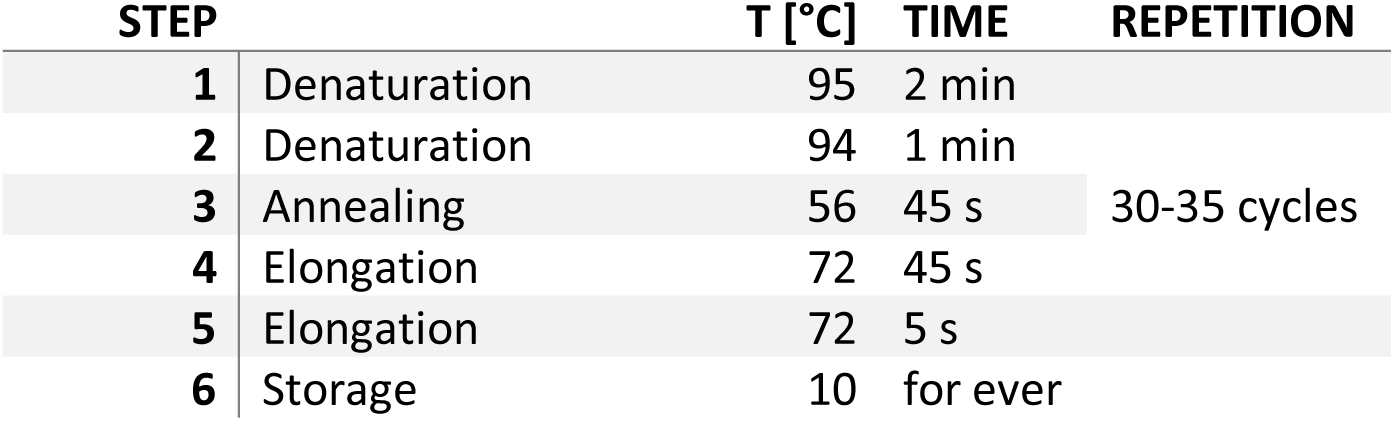

**Text S4.** *Bioinformatics to obtain sex-specific variants based on a male reference genome of A. ruthenus*

Variant calling using all data resulted in a number of 15,682.485 genome-wide variant calls (avg. 8.5 per kbp). Stringent filtering for variants with supporting reads, only present in the female or male pools, resulted in 1,682 female-specific variants and 318 male specific variants (ratio female/male = 5.3). Using different cut-offs for variant filtering, we consistently found the female pool to harbour more sex-specific variants, which hints at a ZW sex determination system. However, the variants did not match known genes related to sex determination. We clustered sex-specific variants by genomic distance and ranked these clusters by variant count. The region with the most female specific variants was HiC_scaffold_4:45,149,338-45,165,057, which contained 27 variants. The second-best cluster consisted of just 6 variants, observed on HiC-scaffold_7:44,514,447-44,516,696 followed by several other hits on HiC_scaffold_4. Looking at male-specific variants, the best region only consisted of 3 variants (HiC_scaffold_4). In the top candidate region, we found a female-specific 15 bp-indel, which we used to design PCR primers, these primers could already sex some of our sexed *A. ruthenus* samples, but not all (details not shown).

**Text S5.** *Corresponding regions in the current A. ruthenus NCBI reference genome*

The current NCBI reference genome (GCF_010645085.1, as of Oct 2020) corresponds to the male genome assembly used for read mapping in our study with a few differences:

A. NCBI assemblies have different sequence identifiers
B. The NCBI assembly has a different gap-size, leading to a shift in coordinates

The changes of sequence identifiers and coordinates are shown in table S1 for the regions mentioned in this study.

---

**Table S1.** Corresponding coordinates of two *A. ruthenus* genome assembly versions.

| Acipenser_ruthenus.ARUT1.2.dna.toplevel.fa | NCBI: GCF_010645085.1 (Oct 2020) |
| --- | --- |
| HiC_scaffold_4: 43,938,061 - 43,987,561 | NC_048326.1: 43,898,461 - 43,947,961 |
| HiC_scaffold_4: 45,149,338 - 45,165,057 | NC_048326.1: 45,108,538 - 45,124,257 |
| HiC_scaffold_7: 44,514,447 - 44,516,696 | NC_048329.1: 44,475,147 - 44,477,396 |

**Table S2.** Data used for phylogenetic tree computation.

| Species | Resource type | Number of contigs/proteins | Resource |
| --- | --- | --- | --- |
| *A. baerii* | transcriptome | 46028 | http://publicsturgeon.sigenae.org/home.html |
| *A. gueldenstaedtii* | transcriptome | 369441 | https://www.ncbi.nlm.nih.gov/Traces/wgs/?page=1&view=tsa&search=acipenser |
| *A. oxyrinchus* | transcriptome | 42114 | http://publicsturgeon.sigenae.org/home.html |
| *A. ruthenus* | genome | 36020 | https://www.ncbi.nlm.nih.gov/genome/?term=txid7906[orgn] |
| *A. schrencki* | transcriptome | 57124 | http://publicsturgeon.sigenae.org/home.html |
| *A. sinensis* | transcriptome | 63077 | http://publicsturgeon.sigenae.org/home.html |
| *A. transmontanus* | transcriptome | 55806 | http://publicsturgeon.sigenae.org/home.html |
| *H. huso* | transcriptome | 548535 | Trinity assembly from mixed organ |
| *L. chalumnae* | genome | 20932 | https://www.ncbi.nlm.nih.gov/genome/?term=coelacanth |

**Figure S1.**
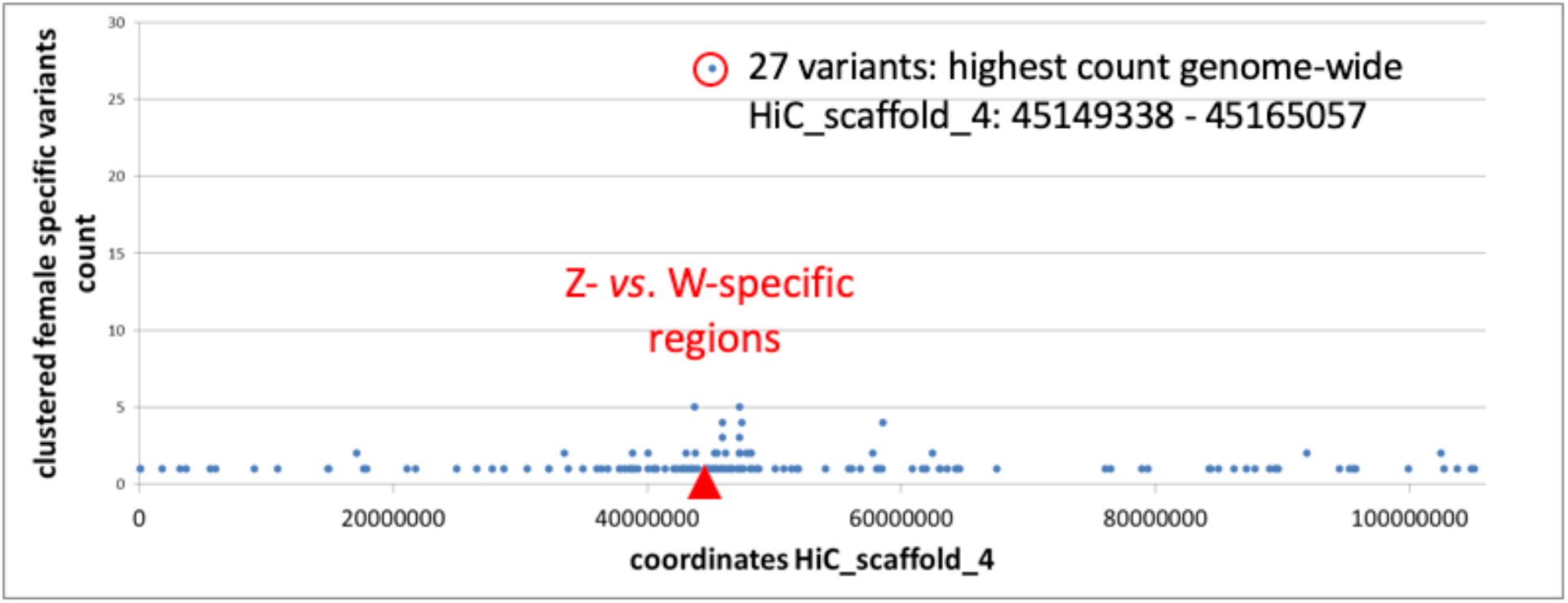
Counts of female-specific variants, clustered according to their distance on HiC_scaffold 4.

**Figure S2.**
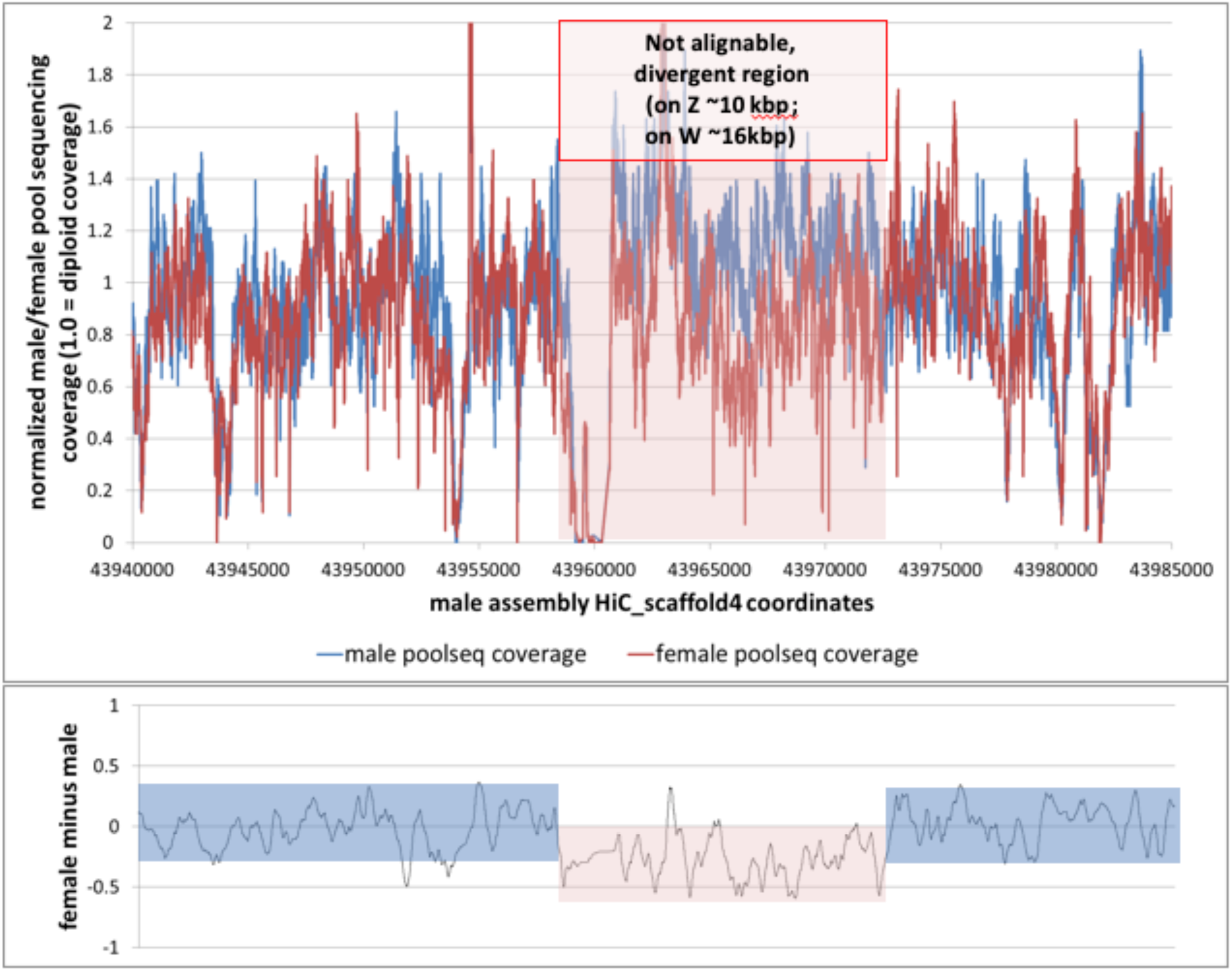
Mapping of pool-seq data of *A. ruthenus* to the male (ZZ) reference genome, female pool-seq data (ZW) shows a ~10 Kb region with reduced coverage. Due to overall coverage fluctuations, it is difficult to reliably detect small loci with haploid coverage differences in genome-wide approaches.

**Figure S3.**
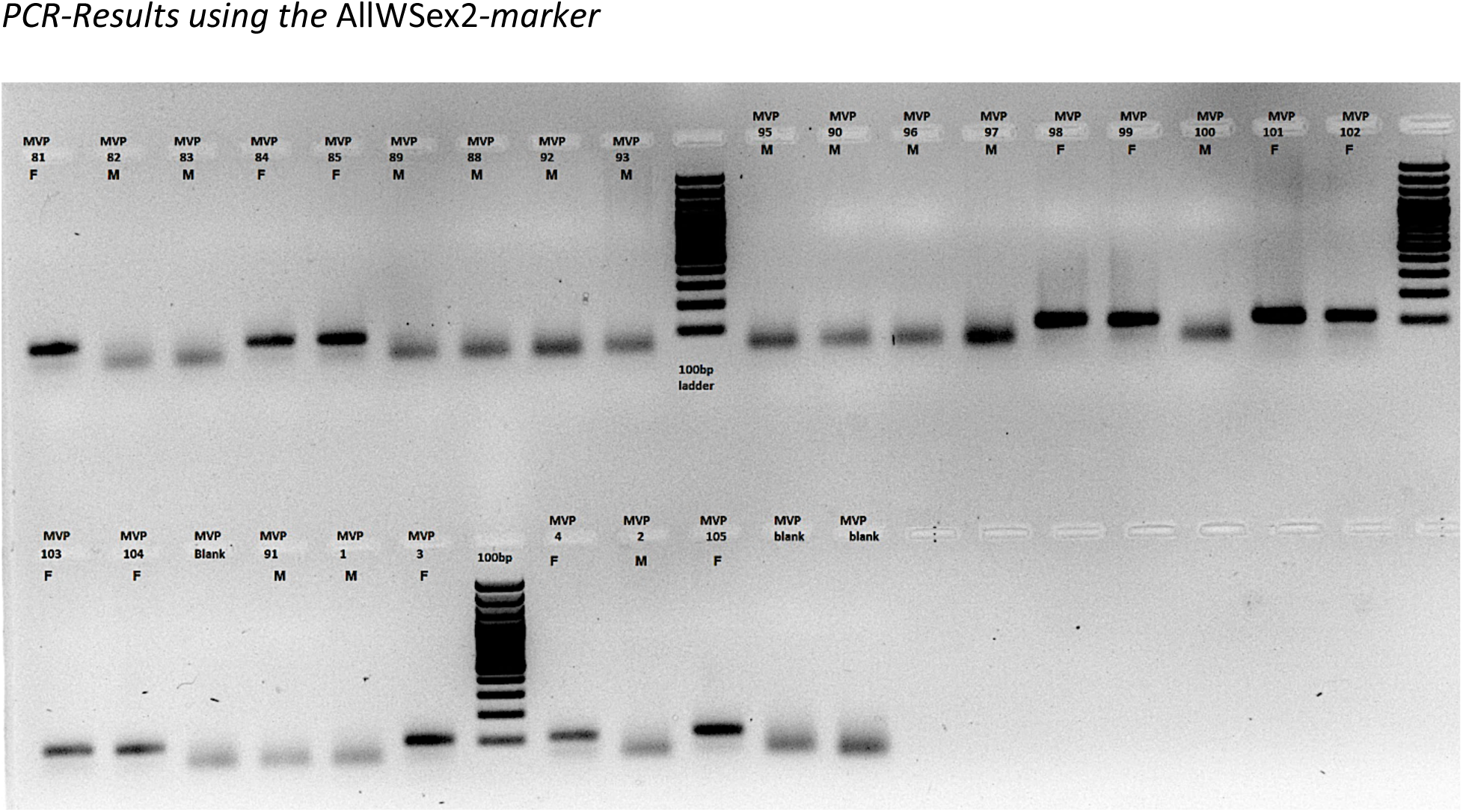
PCR-Result using primers AllWSex2F/R for 26 *A. oxyrinchus* (12 females F, 14 males M) with known phenotypic sex (MVP presents internal sample numbers)

**Figure S4.**
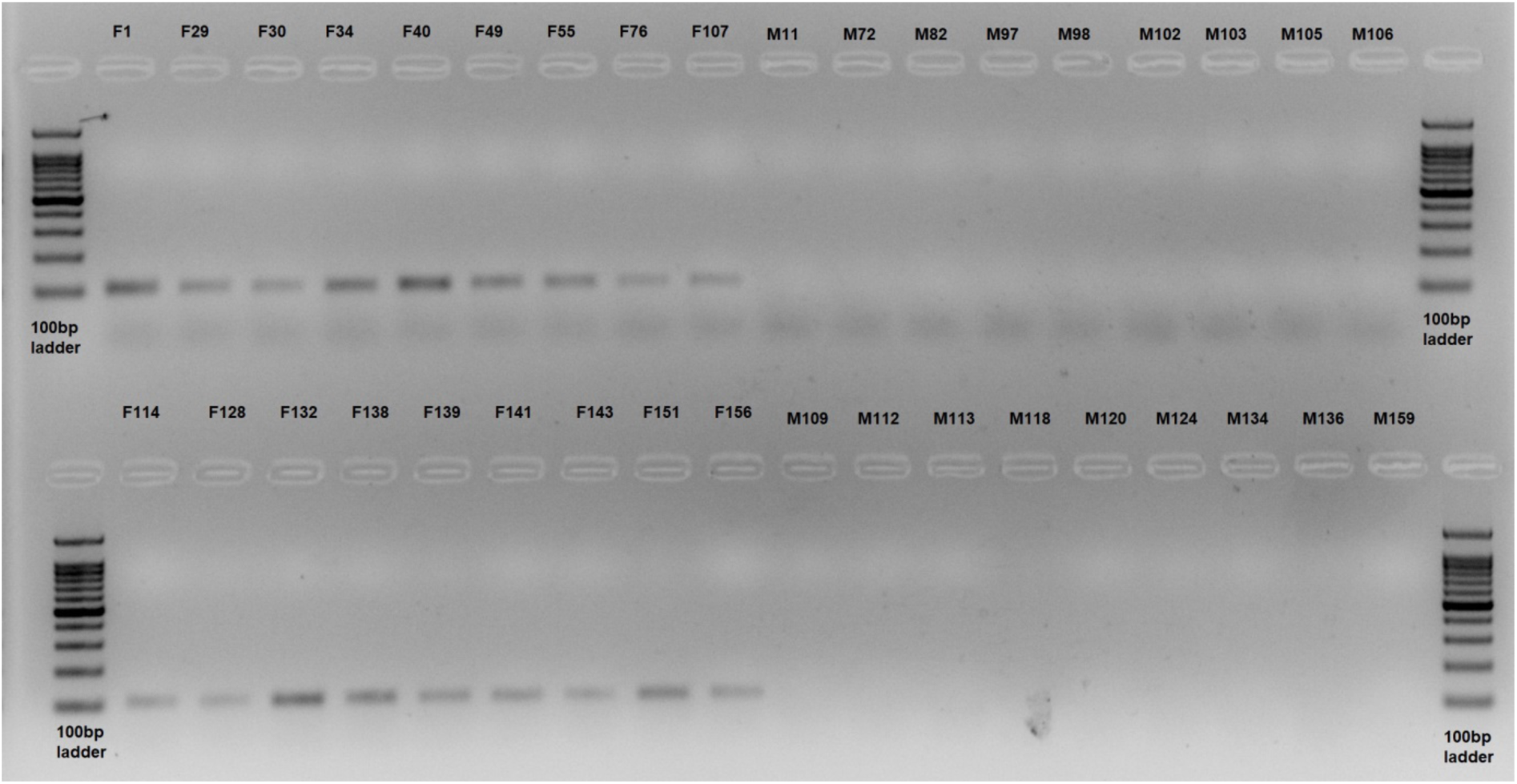
PCR-Result using primers AllWSex2F/R for 36 *A. sturio* (18 females, 18 males) with known phenotypic sex.

**Figure S5.**
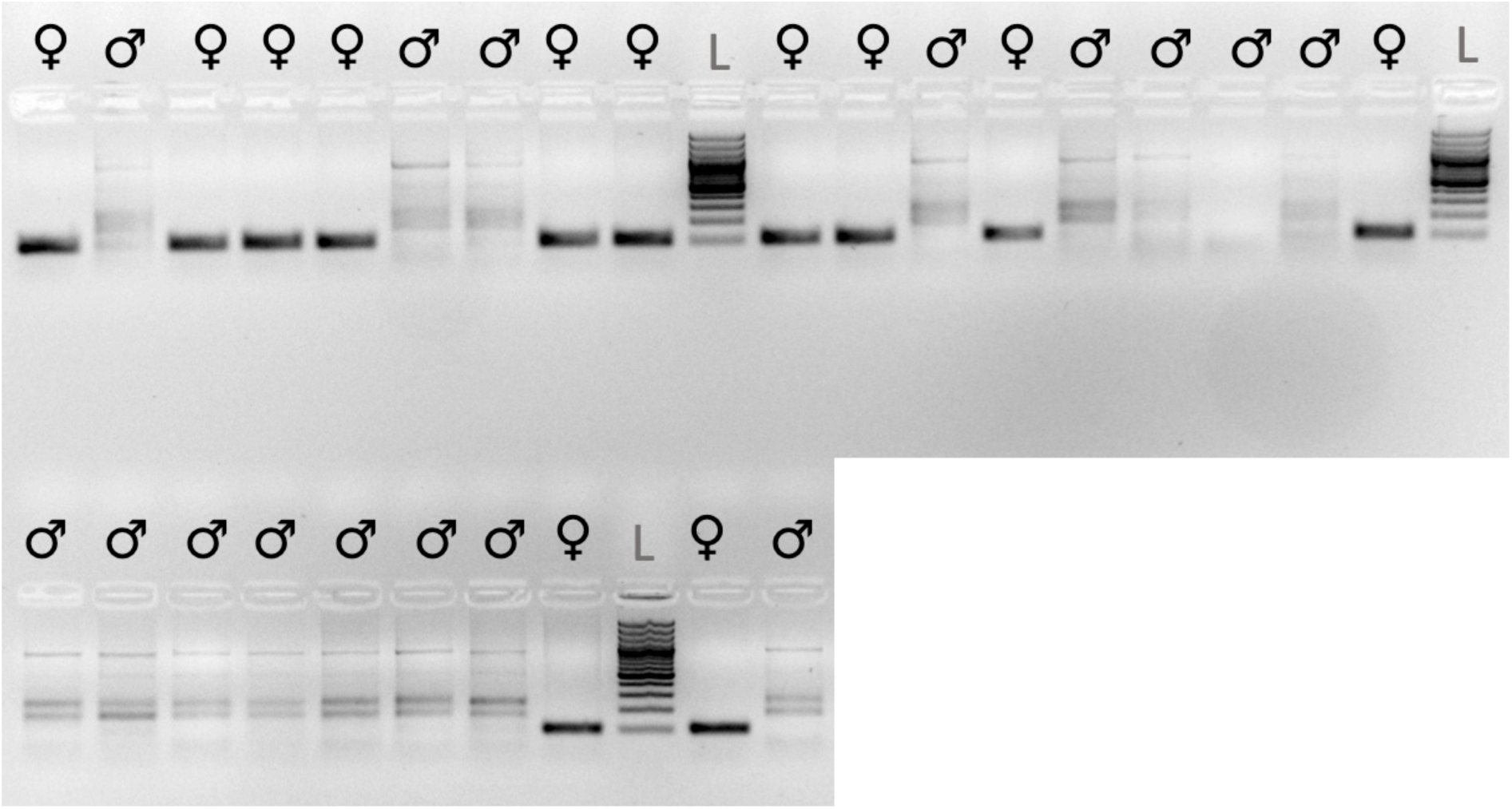
PCR-Result using primers AllWSex2F/R for 28 *Huso huso* (12 females, 16 males) with known phenotypic sex.

**Figure S6.**
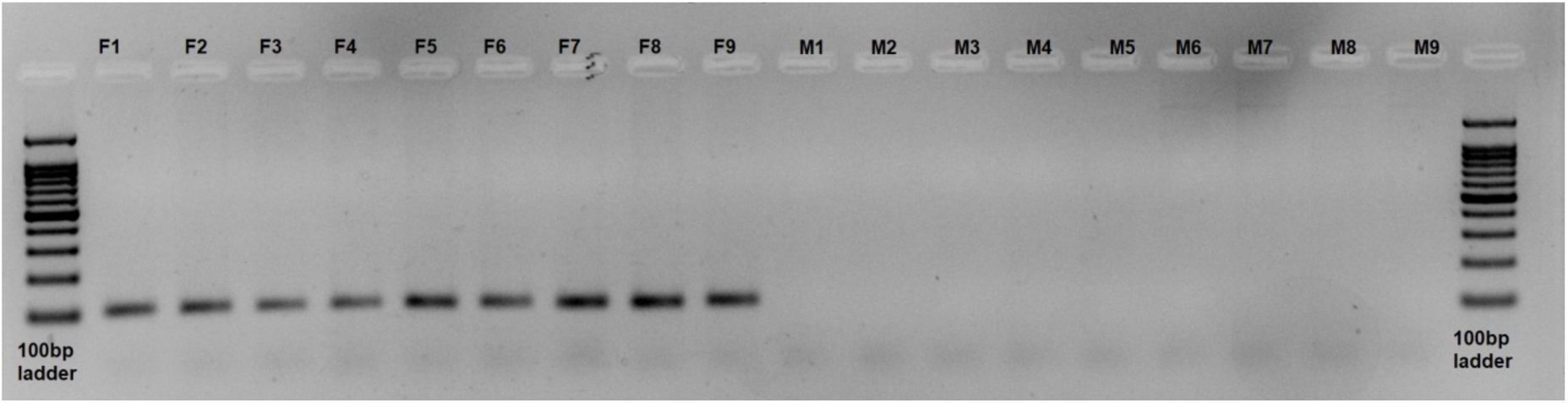
PCR-Result using primers AllWSex2F/R for 18 *A. gueldenstaedtii* (9 females, 9 males) with known phenotypic sex.

**Figure S7.**
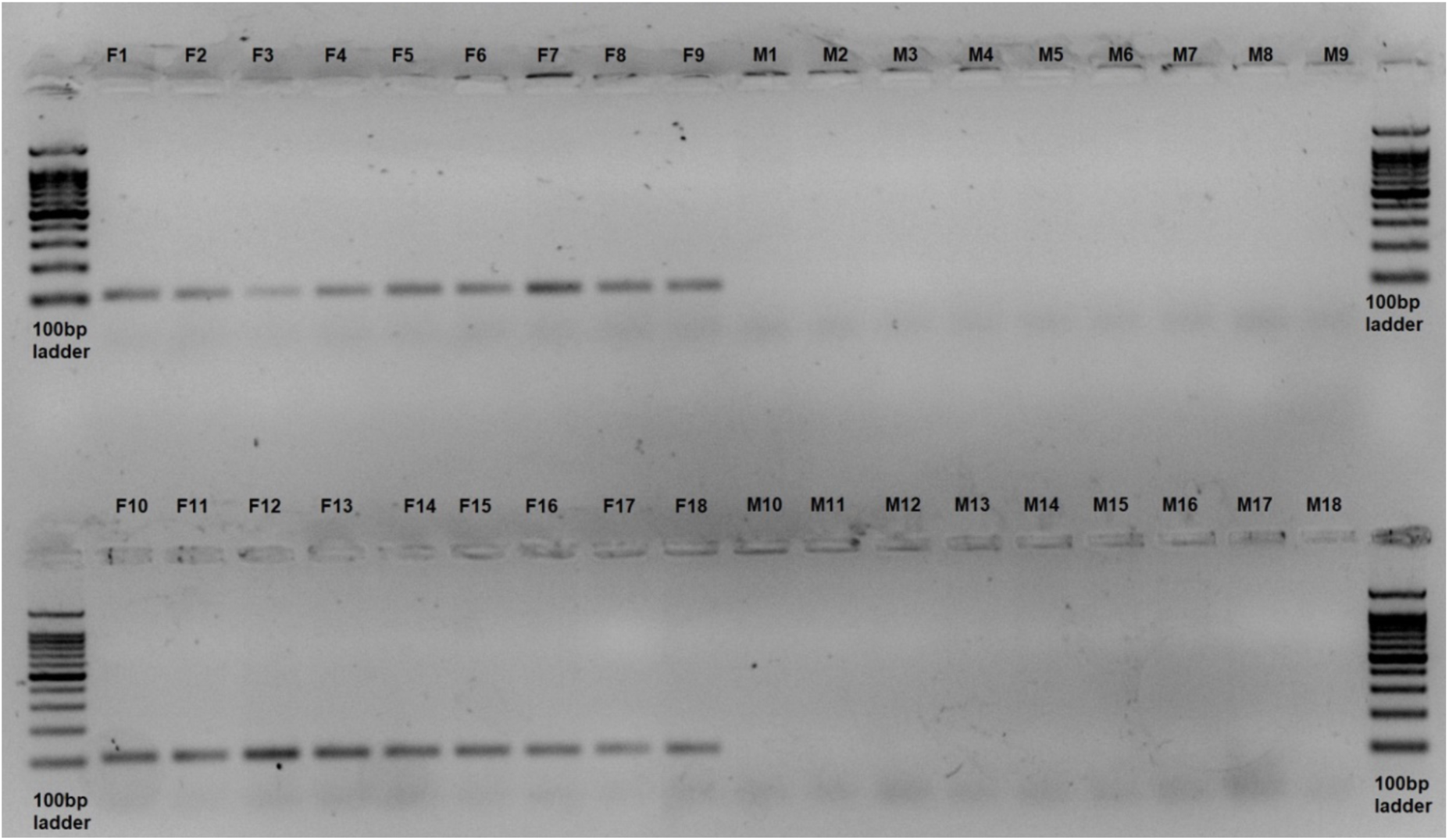
PCR-Result using primers AllWSex2F/R for 18 *A. baerii* (9 females, 9 males) with known phenotypic sex.

**Figure S8.**
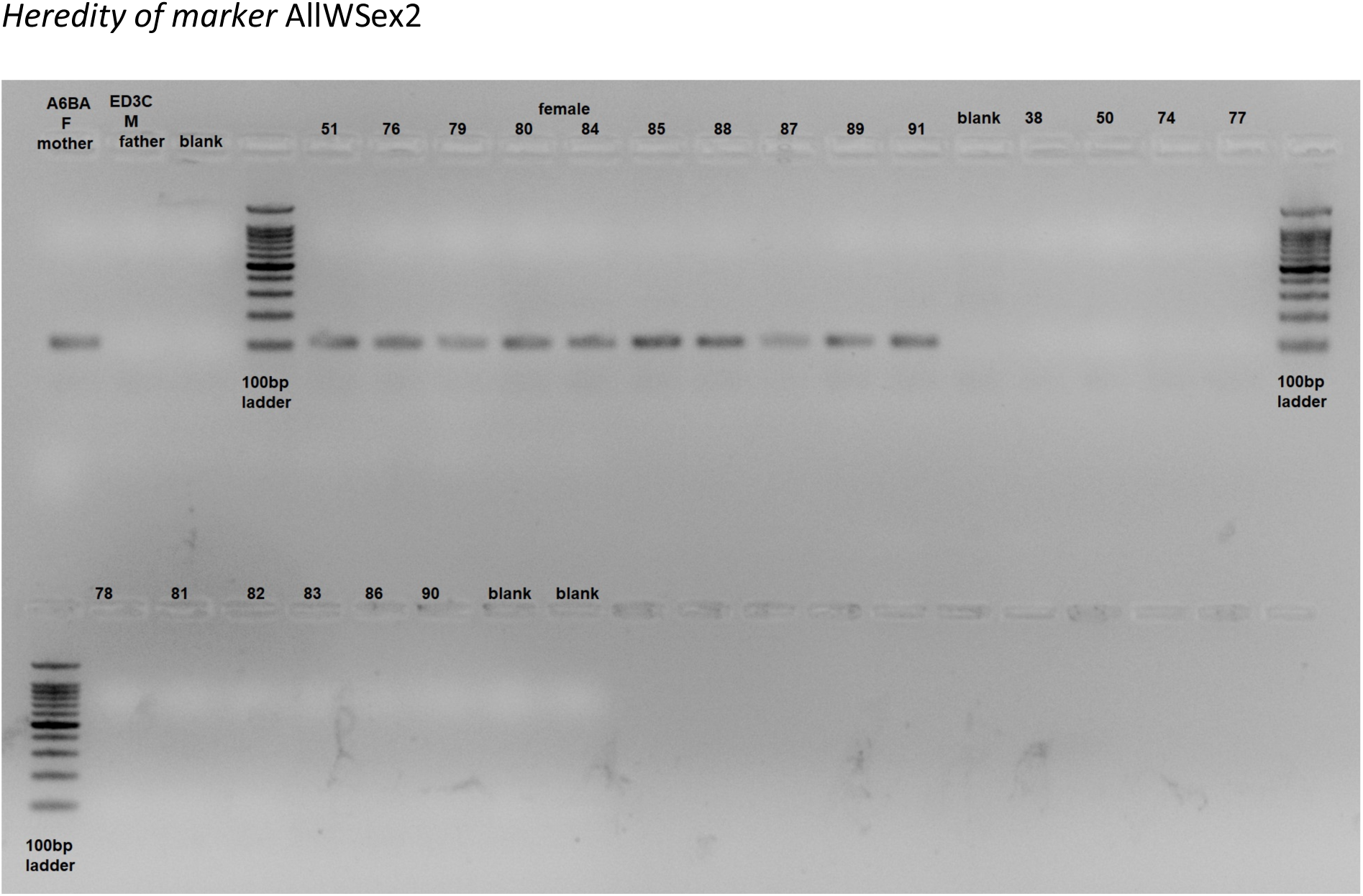
Inheritance of the PCR-marker *AllWSex2* in a genetic family of *A. oxyrinchus*, with mother (A6BA), father (ED3C) and 10 daughters, followed by 10 sons.

**Figure S9.**
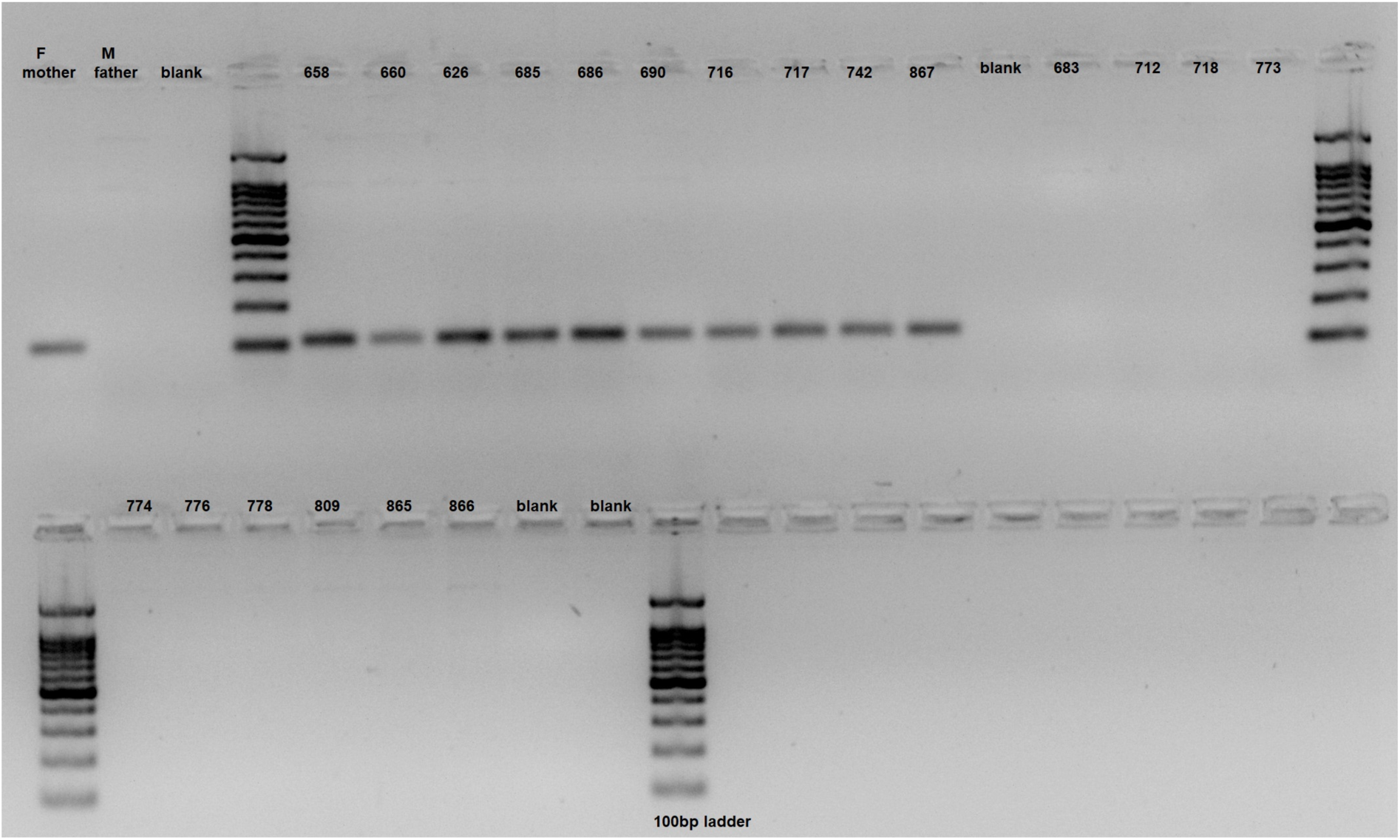
Inheritance of the PCR-marker *AllWSex2* in a genetic family of *H. huso*, with mother (F), father (M) and 10 daughters, followed by 10 sons.

**Figure S10.**
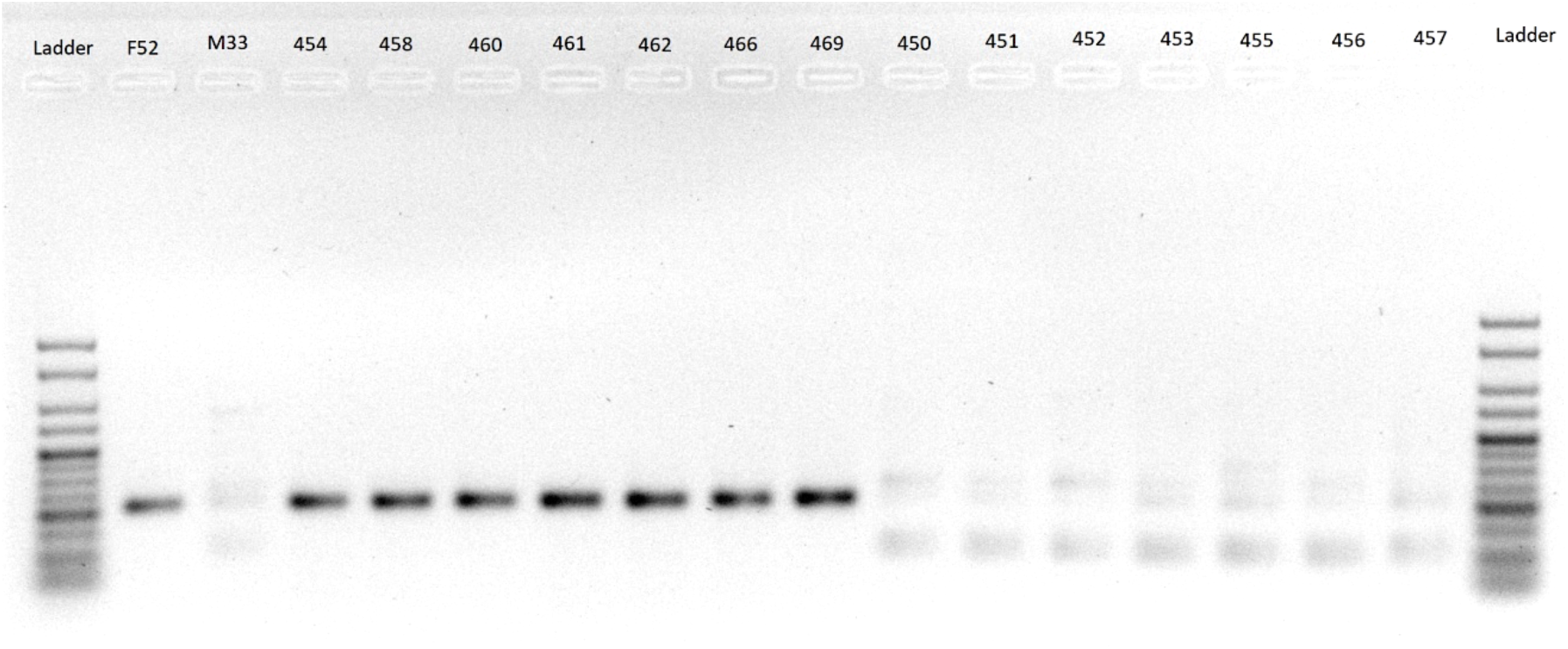
Inheritance of the PCR-marker *AllWSex2* in a genetic family of *A. ruthenus*, with mother (F52), father (M33) and 7 daughters, followed by 7 sons.

**Figure S11.**
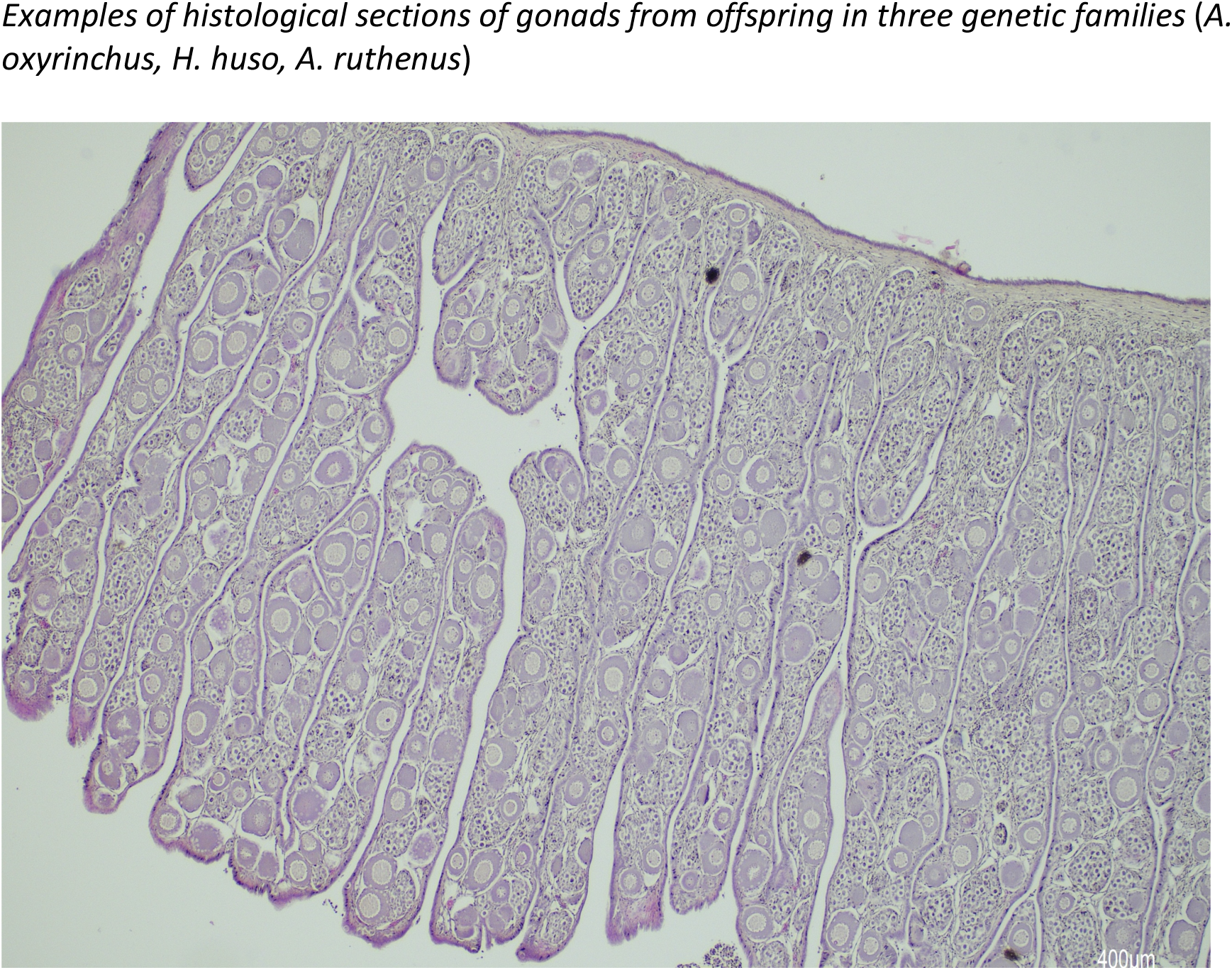
Histology of an *A. oxyrinchus* early female gonad (JG17Ao_F_79_0001.jpg).

**Figure S12.**
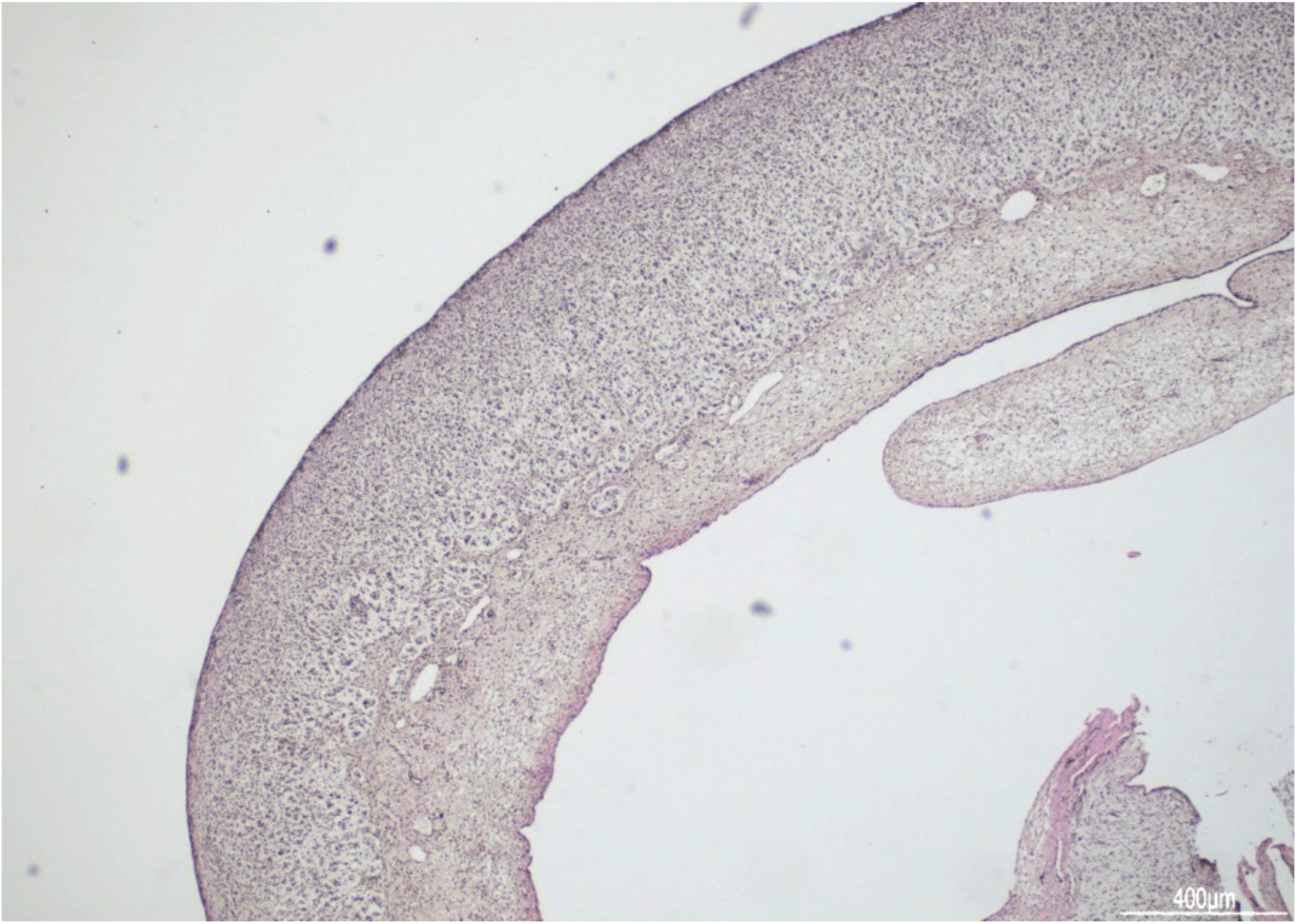
Histology of an *A. oxyrinchus* early male gonad (JG17Ao_M_74_0001.jpg).

**Figure S13.**
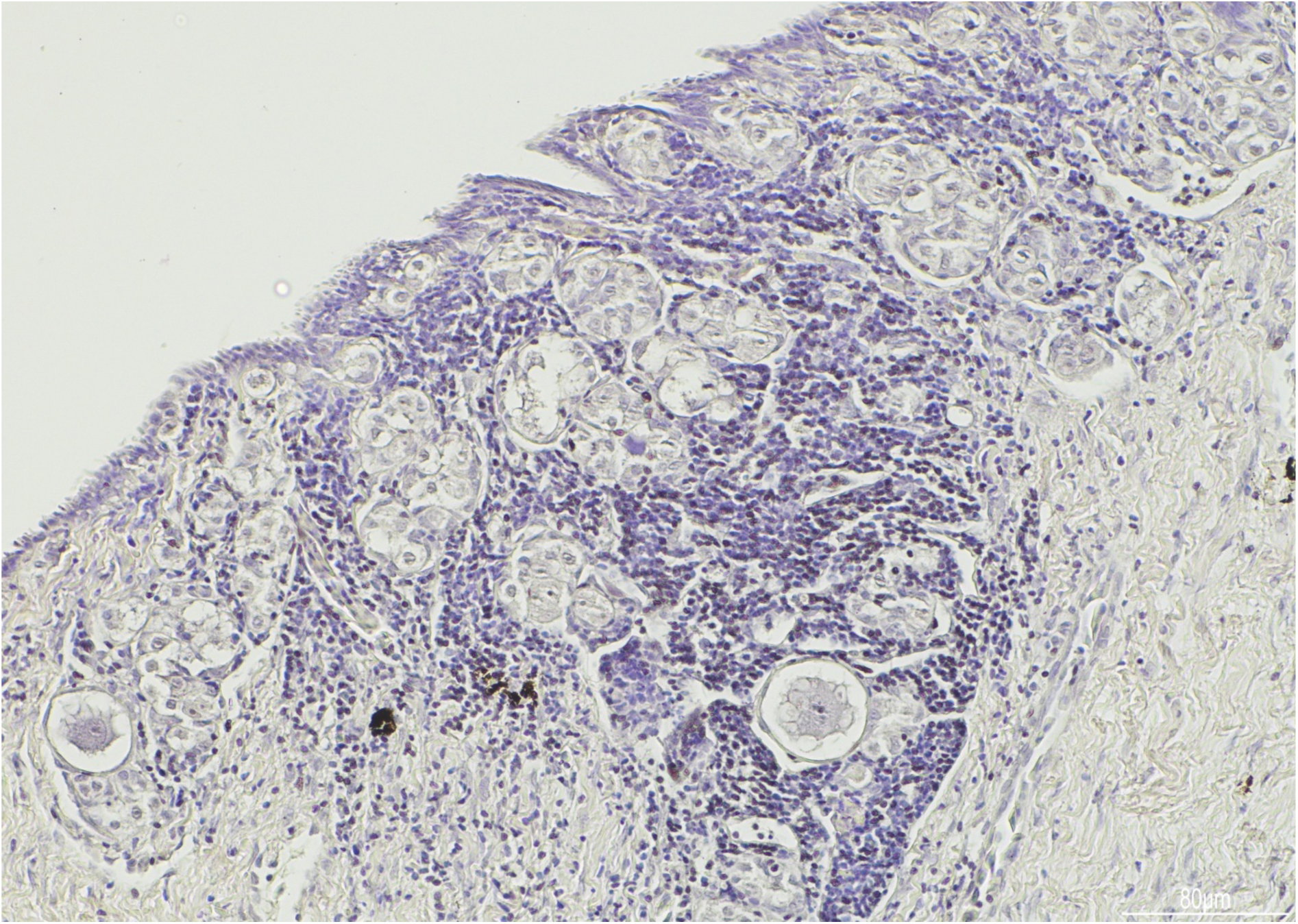
Histology of a *H. huso* early female gonad (Hh_F_H446R626_0006.jpg).

**Figure S14.**
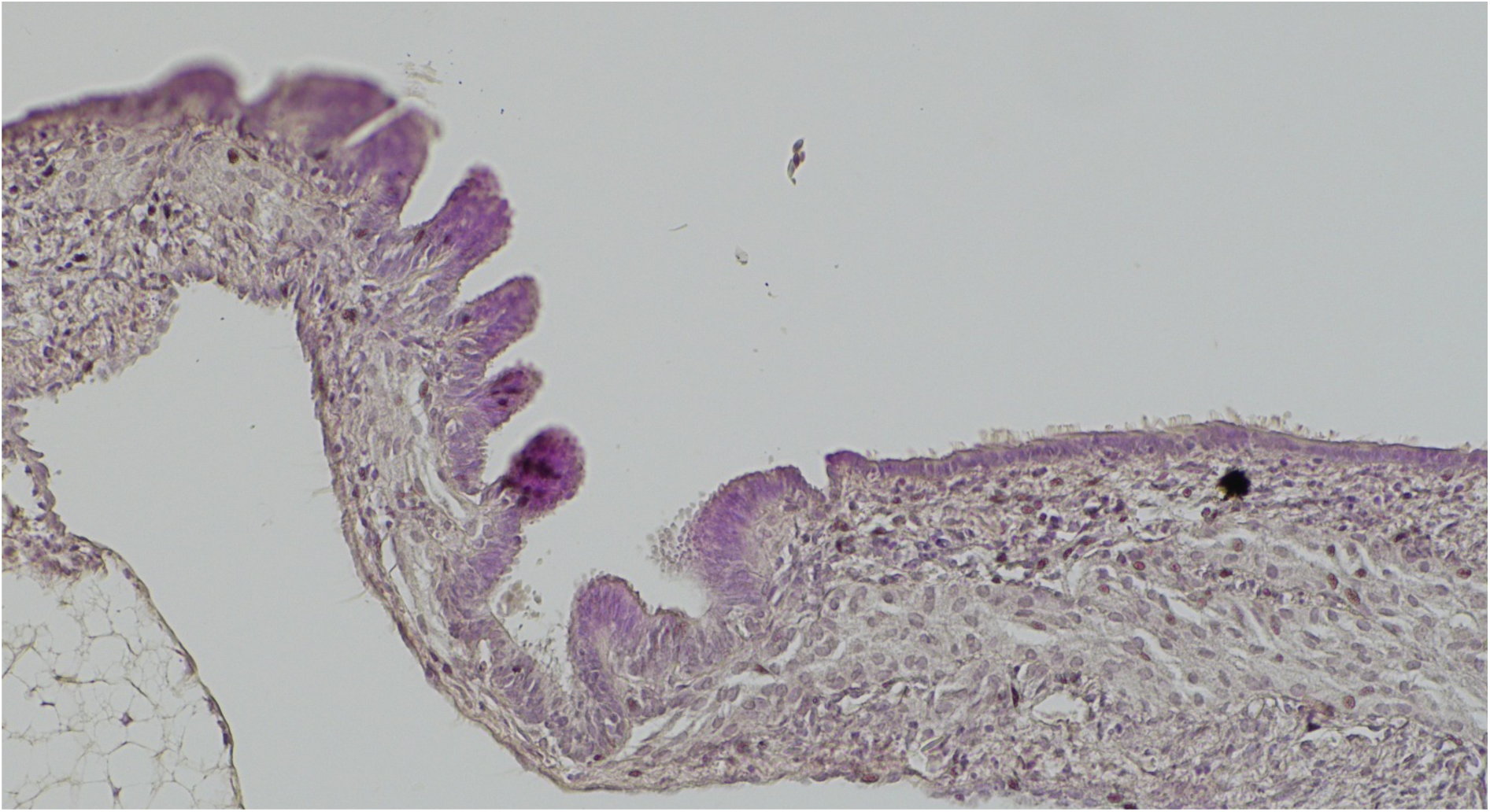
Histology of a *H. huso* early male gonad (Hh_M_H593R773_0001.jpg).

**Figure S15.**
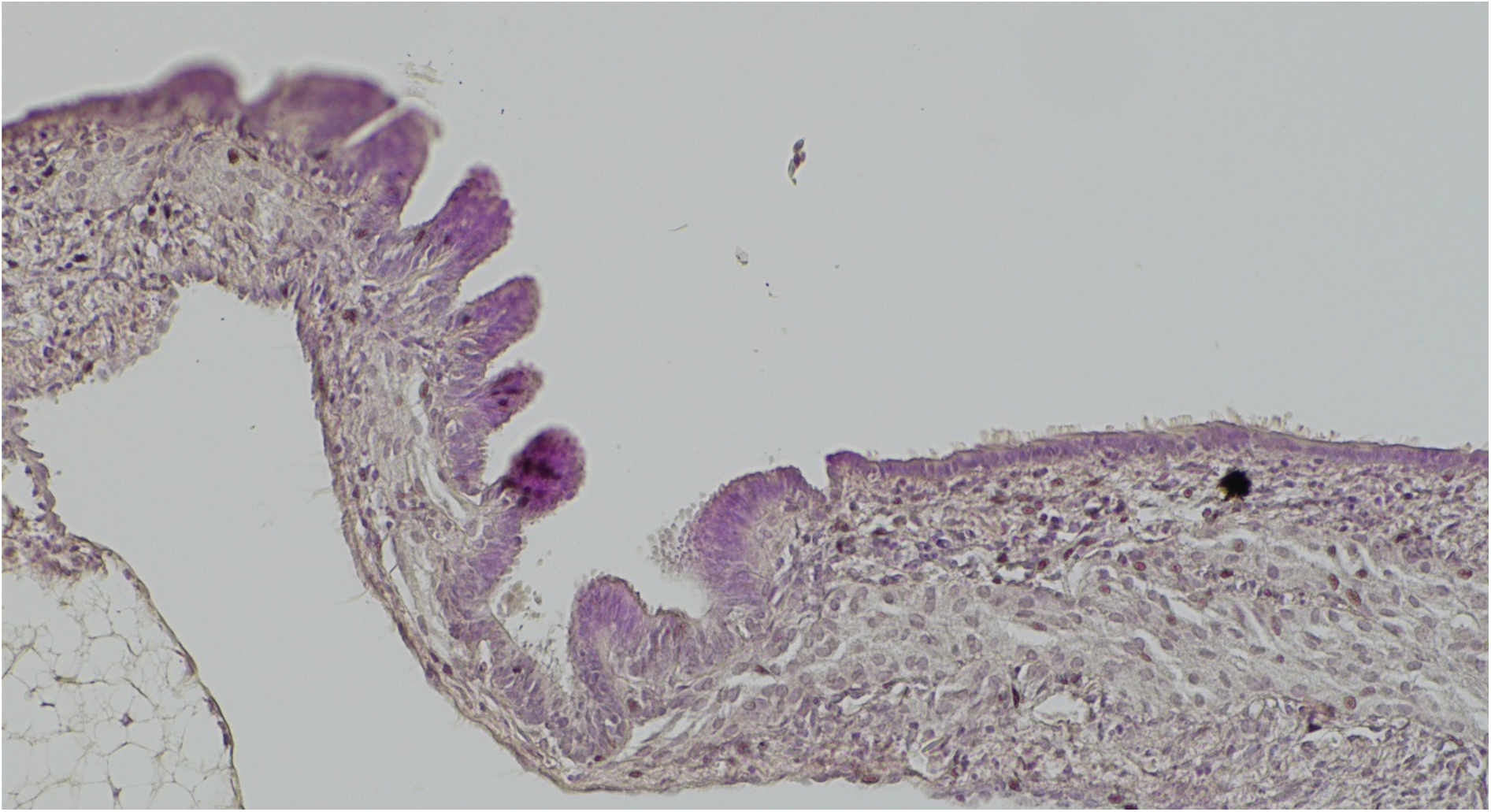
Histology of an *A. ruthenus* early female gonad (AruthX3_F_458_0018.jpg).

**Figure S16.**
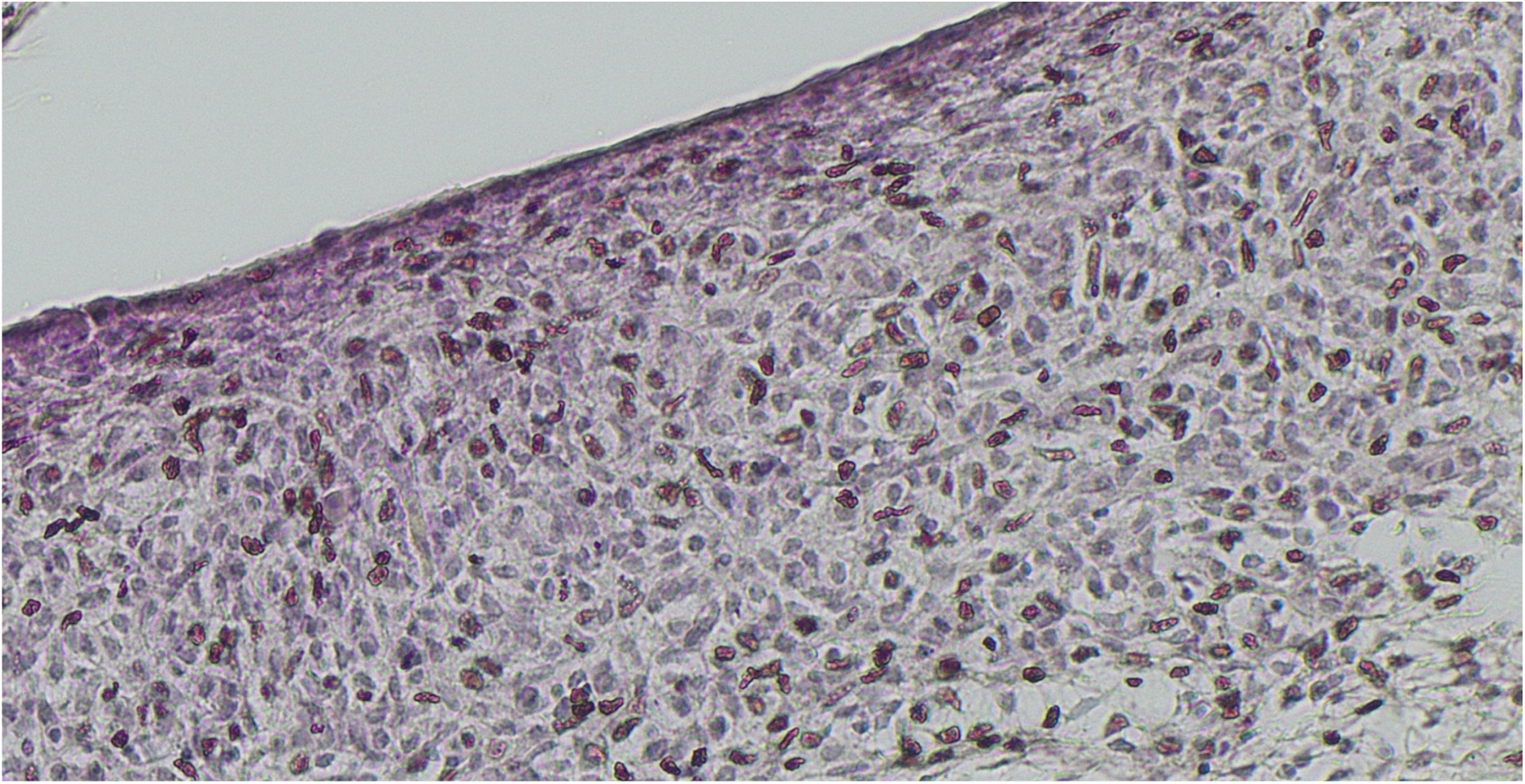
Histology of an *A. ruthenus* early male gonad (AruthX3_M_457.jpg).

